# AAV-Delivered Anti-PC-OxPL Antibody Fragments: A Novel Therapeutic Approach to Target ALS

**DOI:** 10.1101/2025.01.22.634350

**Authors:** Andreia Gomes-Duarte, Jamie K. Wong, Svetlana Pasteuning-Vuhman, Wouter Pos, Rob Haselberg, Marina Sogorb-Gonzalez, Sofia Pascoal, Roland van Dijk, Farangis Sharifi, Martin Giera, Pavlina S. Konstantinova, Sotirios Tsimikas, Saud A. Sadiq, Sander van Deventer

## Abstract

Amyotrophic lateral sclerosis (ALS) is characterized by the progressive loss of motor neurons and premature death. The limited understanding of the mechanisms underlying selective motor neuron death has significantly hindered the development of disease-modifying treatments. Ferroptosis, a form of cell death dependent on iron accumulation, has been implicated in the selective degeneration of motor neurons in ALS. Oxidized phosphatidylcholines (PC-OxPL) have been identified as key effectors in the pathophysiological processes associated with this pathway. Preclinical observations revealed a distinct PC-OxPL profile in the cerebrospinal fluid (CSF) of sporadic ALS (sALS) patients and identified apolipoprotein E (APOE) particles as the primary carriers of PC-OxPL in the CSF. Furthermore, in ALS brain and spinal cord tissue sections, PC-OxPL was found to be predominantly associated with motor neurons. Exposure of iPSC-derived motor neurons to PC-OxPL led to transcriptomic changes in genes known to be linked to ALS, as well as the induction of significant TDP-43 pathology and motor neuron death. To counter this, we developed a single-chain antibody fragment (scFv) encoded by an AAV-delivered DNA construct that specifically targets PC-OxPL neoepitopes (PC-OxPL-VecTab®). PC-OxPL-VecTab® effectively neutralized PC-OxPL-induced neurotoxicity and TDP-43 aggregation in motor neurons, while preventing motor neuron death and deficits in a sALS CSF mouse model. When administered intrathecally to minipigs, PC-OxPL-VecTab® was distributed to both upper and lower motor neurons and expressed at levels predicted to be therapeutically effective. Our work identifies PC- OxPL as a critical pathological factor and a key inducer of TDP-43 pathology in ALS, providing the foundation for a novel therapeutic intervention modality for patients with sALS. Furthermore, it offers the exciting potential to be expanded to diseases characterized by PC-OxPL neurotoxicity.

## Introduction

Amyotrophic lateral sclerosis (ALS) is a degenerative neuromuscular disease characterized by the progressive degeneration of motor neurons in the brain and spinal cord. Most patients die within three to five years of -diagnosis, typically due to respiratory failure ^1,2^. Current treatments largely focus on symptom management ^3^. While a small subset of ALS cases is linked to mutations in genes such as SOD1, C9orf72, or FUS, the majority are classified as sporadic ALS (sALS). TDP-43 pathology or proteinopathy, present in 97% of all ALS patients, involves the mislocalization of TDP-43 from the nucleus to the cytoplasm, leading to the formation of toxic aggregates ^4^. TDP-43 is essential for chromatin remodeling, RNA metabolism, proteostasis, stress granule formation, and mitochondrial function and dysfunction of TDP-43 has been shown to impact motor health negatively ^4,5^ .

Several mechanisms of programmed cell death including apoptosis, necroptosis and ferroptosis have been proposed to cause motor neuron death in ALS ^6,7^. However, evidence does not support apoptosis as a key driver, and a recent clinical trial targeting RIPK1, a critical necroptosis mediator, failed to show therapeutic benefits in ALS patients (Phase 2 HIMALAYA; SAR443820). Conversely, ferroptosis - a form of cell death driven by iron-dependent reactive oxygen species (ROS) formation - has been shown to selectively affect motor neurons in ALS ^8^. Glutathione peroxidase 4 (GPX4), the main intracellular inhibitor of ferroptosis, plays a crucial role, and GPX4-deficient mice exhibit motor neuron-specific degeneration ^6,8^. A distinct feature of ferroptosis is the extensive oxidation of poly-unsaturated fatty acids (PUFA) in comparison to other forms of cell death ^9^. In ALS, motor neurons exhibit increased glycerophospholipid metabolism compared to unaffected ocular motor neurons ^10^. In particular, elevated levels of phosphatidylcholines (PC) containing native phospholipids, precursors of PC-containing oxidized phospholipids (PC-OxPL), in cerebrospinal fluid (CSF) correlate with disease progression ^10,11^. Furthermore, histological studies have detected PC-OxPL accumulation in upper motor neurons of ALS patients, and high systemic levels of oxidized low-density lipoproteins (OxLDL), enriched in PC-OxPL, have been associated with rapid ALS progression ^12,13^.

Mice possess natural IgM E06 antibodies at birth, which specifically recognize the PC headgroup of oxidized but not native phospholipids ^14^. Transgenic mice overexpressing a single chain variant of E06 have been documented to ameliorate a variety of pathogenic states, including atherosclerosis, hepatic steatosis, ischemia reperfusion injury in the heart and liver, improvements in bone density and nociceptive pain syndromes ^15–17^. The central nervous system (CNS) is not known to contain these innate antibodies, and growing evidence suggests PC-OxPL are linked to neurotoxicity in neurodegenerative diseases ^18,19^.

This study investigated the mechanisms underlying PC-OxPL-induced neurotoxicity in ALS and evaluated the therapeutic potential of neutralizing PC-OxPL using an AAV-delivered PC-OxPL- VecTab®. Our findings demonstrate that PC-OxPL species are increased in the brain of ALS patients with APOE identified as the primary PC-OxPL carrier in the CNS and upregulated in ALS. We show that PC-OxPL exposure in wild-type (wt) and ALS (TDP-43^M337V^) motor neurons induced ALS-relevant gene expression, triggered TDP-43 aggregation, and resulted in motor neuron toxicity. Remarkably, PC-OxPL-VecTab® effectively neutralized PC-OxPL toxicity *in vitro*, while preventing the pathology caused by sALS CSF transfer into wt mice, such as TDP-43 mislocalization, motor neuron death, and motor deficits. Furthermore, safety and biodistribution studies in mini pigs confirmed the therapeutic applicability of PC-OxPL-VecTab® in humans. Our results highlight the accumulation of PC-OxPL as a key pathological mechanism in ALS, suggesting that targeting PC-OxPL neurotoxicity is a promising strategy for ALS therapies.

## Materials and Methods

### Human samples and tissue characterization

#### Sample collection

Details on sample selection, demographic data, and disease characteristics can be found **in Supplementary Table S1**.

#### Immunohistochemistry and imaging of human tissue samples

FFPE tissue samples were obtained from the Netherlands Brain Bank (NBB). FFPE tissues were sectioned with Leica Biosystems Cryostat (Leica RM2255). Samples were trimmed until the entire section profile was visible. Sectioning continued until 30 consecutive 7 μm sections were retained.

For deparaffinization and rehydration, sections were incubated in Tissue Clear for 10 minutes (min). Then, sections were incubated in Tissue Clear/100 % Ethanol (ETOH) (1:1) for five min, 100% ETOH for two min, 96% ETOH for two min, 70% ETOH for two min, and 50% ETOH for two min. Finally, the sections were washed in phosphate-buffered saline (PBS) for five minutes and incubated with ddH_2_O for two minutes.

A Tris-based buffer was prepared by adding 2.5 ml of Antigen Unmasking Solution (Tris-based) to 250 ml of ddH2O and pH-adjusting to 9.0. Slides were incubated in a pressure cooker at 95°C for 40 min and then cooled for 20 min. Sections were washed two times in PBS for three min.

Sections were encircled with a grease pen, and slides were sorted into a damp chamber. Sections were incubated with 1x TrueBlack (Biotinum, cat. No. 23007) in 70% ETOH for 30 seconds (s) (100-150 μL per section) and washed two times for five min in PBS. Sections were blocked for 30 min with 10% donkey-blocking serum solution at room temperature (RT). The blocking buffer was wiped away, and sections were incubated with primary antibodies overnight at RT. Sections were washed three times for five min in PBS and incubated with secondary antibodies for one hour (h) at RT. Sections were washed twice for five min in PBS, followed by staining for 20 min with 4′,6-diamidino-2-phenylindole (DAPI) solution. Sections were washed twice for five min in ddH_2_O and finally embedded in Mowiol (stored in a briefcase). Antibody details and experimental conditions are listed in detail in **Supplementary Table S2**.

For imaging, Zeiss Axio Scan Z1 was used with 10 x lens, Plan Apochromatic (NA 0.45) or Fluar (NA 0.5) air.

#### Image processing and analysis

Images were initially obtained in .czi format. For image export, the original 16-bit format was converted to an 8-bit format. Images were processed using the following software: Zen Blue Edition (v 3.1 or higher), Image Pro Premier (v 9.3 or higher), and Fiji/Image J (v 1.54f).

Line profile was generated in Zen lite using distance and intensity measurements for the channels of interest.

#### APOE enzyme-linked immunosorbent assay (ELISA)

Samples were assayed using the RayBio® Human APOE ELISA Kit. The assay was run following the manufacturer’s instructions; kit lot 0301240061 was used for the analysis. Samples were diluted at 1:3000 and 1:500 for plasma and CSF, respectively. In brief, after incubation with APOE standard or plasma/CSF samples the wells were sequentially treated with biotinylated anti-APOE, streptavidin and TMB One-step substrate reagent with buffer washes in between each step. After adding the TMB substrate, the plate was incubated for 30 min at RT under gentle shaking (450 rpm), after which 50 μL stop solution was added. Signals were acquired after five min by using the EnVision Reader at 450 nm. To determine APOE concentrations, standard curve signals were log-log transformed and fitted by a 5- parameter logistic fitting model by using the GraphPad Prism V 9.5.1 Software. The back-calculated analyte amount was adjusted for the dilution factor to determine its concentration in the CSF samples. Relative APOE amount in ALS samples was calculated as a percentage concerning the average amount measured in healthy controls (HC) (defined as 100%).

#### Quantitation of PC-OxPL in human CSF on lipoproteins

Established and novel chemiluminescent enzyme linked assays were used to quantitate PC-OxPL on APOB-100 (APOB), APO(a), APOE, APOC-III and APOA-I in plasma. OxPL-APOB, OxPL-APO(a) and OxPL-APOAI have been validated and previously described in detail ^17,20,21^. Briefly, antibodies MB47 binding to APOB, LPA4 binding to APO(a) and sheep anti-human APOA-I (The Binding Site, Birmingham, UK) are plated overnight at µg/ml to microtiter well plates, plasma is added at 1:50 dilution and OxPL-APOB and OxPL-APO(a) are detected by b-E06 IgM and measured by chemiluminescence as relative light units per 100 milliseconds (RLU). A standard curve of PC equivalents using a linear range of PC-BSA is then used to convert RLU to nmol/L PC-OxPL which is additionally adjusted for the plasma dilution. To detect OxPL-APOE and OxPL-APOC-III, microtiter well plates were coated with polyclonal rabbit anti-human APOE (ThermoFisher Scientific, Waltham, MA) that recognizes all three APOE isoforms at 5 µg/ml and a rabbit monoclonal antibody to APOC-III (Abgent, San Diego, CA) at 2 µg/ml after documenting these concentrations were saturating. Plasma was added at 1:50 dilution and OxPL-APOE and OxPL-APOC-III were detected and reported in nmol/L as above. For CSF, because the concentration of analytes is lower, a 1:10 dilution of CSF was used and OxPL-APOB, OxPL-APO(a), OxPL-APOE, OxPL-APOC-III and OxPL-APOA-I were measured and reported after adjusting for appropriate dilution in a similar fashion to the plasma assays.

#### Targeted lipidomics analysis of human CSF and plasma

A targeted LC-MS assay covering 31 PC-OxPL species and their precursors ^22^ was used to quantify their levels in the plasma of ALS patients and HC. The complete list of lipid species monitored and detected can be found in **Supplementary Table S3**. In short, lipids were extracted using a modified methyl tert-butyl ether-based protocol from 25 µL of CSF or 25 µL of plasma after the addition of 20 µL of an internal standard mix (100 ng/mL DNPC and d5-PC 17:0/22:4 in methanol). After drying and reconstitution, samples were analyzed on a Shimadzu Nexera LC40 system coupled to a Sciex 6500+ QTrap mass spectrometer, following published protocols ^22^.

## In vitro Experiments

### Human iPSC-derived cultures

#### Axol motor neuron progenitors (MNP)

Healthy human iPSC-derived motor neuron progenitors (MNPs) (ax0078, Axol Bioscience) were purchased and used according to the manufacturer’s protocol. In brief, a T25 culture flask (734-2004, VWR) was coated with 0.1 mg/mL PDL (A38904-01-Gibco) for 2 h at 37°C followed by three times wash with PBS (90-094, Gibco). Next, the culture flask was coated with 20 µg/mL Surebond (ax0041, Axol Bioscience) overnight at 37°C. Cryopreserved MNPs were thawed and resuspended in recovery medium (ax0071, Axol Bioscience) supplemented with 0.1 µM retinoic acid (R2625-50MG, Sigma) and 10 µM Y-27632 (10-2301, Focus Biomolecules). Cells were plated at a density of 1-1.5E5 cells/cm² in the PDL/Surebond-coated T25 flask and cultured in a humidified incubator with 5% CO_2_ at 37°C. The day after seeding, the medium was replaced completely with recovery medium without Y-27632. Full medium changes were performed every 2-3 days. At a confluence of 70%, MNPs were passed and expanded. For the dissociation, cells were incubated with Accutase (A11105-01, Gibco) and monitored until >90% of cells were detached. Cells were seeded in motor neuron maintenance medium completed with 0.2 µM Compound E (ab142164, Abcam), 0.5 µM retinoic acid, 5 ng/mL BDNF (78133, Stem Cell Technologies), 10 ng/mL CNTF (78170, Stem Cell Technologies), and 10 µM Y-27632 in a 96- well plate (165305, Thermo Fisher Scientific). The next day, the medium was replaced with complete motor neuron maintenance medium without Y-27632, and then half of the medium was replaced every two to three days.

#### Wt and ALS-derived motor neurons and astrocytes

Human iPSC-derived cell lines (iCell Motor Neurons and Astrocytes) were purchased from Fujifilm Cellular Dynamics International (FCDI, Madison, WI). Frozen vials were thawed and seeded according to the manufacturer’s instructions at a recommended density of 100.000 live cells/cm^2^. iCell Motor neurons were cultured in a volume of 500 or 100 µL on a 12- (3513, Corning) or 96-well plate (165305, Thermo Fisher Scientific), respectively.

#### Co-culture

The co-culture seeding medium was prepared by combining DMEM/F12 (11320033, Gibco), Neurobasal medium (21103049, Gibco), B27 Supplement (17504044S, Gibco), N2 Supplement (17502048, Gibco), GlutaMax (35050061, Gibco), human recombinant BDNF (78133, StemCell Technologies), human recombinant GDNF (78139, StemCell Technologies), human recombinant TFG- β1 (100-21C-B, PeproTech), Astrocyte Supplement (BrainXell) and Neuronal Seeding Supplement (BrainXell). iPS- derived motor neurons (BX-0100, BrainXell) and spinal astrocytes (BX-0650, BrainXell) were thawed and seeded at 20.000/5000 cells per well of a 96-well plate (165305, Thermo Fisher Scientific). On day one, all the medium was replaced with fresh medium. On day 4, an equal volume of fresh medium was added, and medium was refreshed every three days. From day 7 onward, medium was added without astrocyte- and day 4 supplement. All cultures were kept at 37°C in a 5% (v/v) CO_2_ atmosphere incubator over the course of the experiments.

#### PSPC and PC-OxPL treatment

*1-palmitoyl-2-(9-oxononanayl)-phosphocholine* (PONPC) (870605P-1MG, Avanti Polar Lipids), *1- palmitoyl-2-azelaoyl-sn-glycero-3-phosphocholine* (PAzPC) (670600P-1MG, Avanti Polar Lipids), and *1-hexadecanoyl-2-octadecanoyl-sn-glycero-3-phosphocholine* (PSPC) (850456P-25MG, Avanti Polar Lipids) were dissolved in 100% ETOH (437433T, VWR) and heated to 35°C followed by sonication (37 kHz) for 30-60 s. Solutions were stored at -150°C for further use in experiments. Fresh PSPC and PC-OxPL solutions were prepared before the start of the experiment and diluted in complete motor neuron maintenance medium to 2x the final concentration. For treatment, half of the medium was removed and replaced with 2x concentrated PSPC or PC-OxPL medium to achieve the final concentration. Cells were treated with concentrations ranging from 25-100 µM and incubated for the assigned time (24 hours or 24 h followed by 24 h washout, depending on the readout). 0.3% EtOH was used as a vehicle control. All medium was removed and replaced with complete motor neuron maintenance medium for the washout experiment.

An initial assessment of neuronal culture health following PSPC and PC-OxPL treatment was performed using live imaging and Calcein AM (Invitrogen) cell-permanent dye. Briefly, a Calcein working solution mix (1:500 in PBS) was prepared and pre-warmed to 37°C before use. At designated time points of analysis post PSPC and PC-OxPL treatment, the medium was removed from the cells and replaced with a working solution mix for 30 min at 37 °C. Finally, the working solution mix was removed, and the cell cultures were incubated with pre-warmed 0.2% BSA in PBS solution, followed by live cell imaging.

#### Generation and transduction of AAV5.2-Control

Mouse PC-OxPL-VecTab® sequences were identical to a previously reported design^23^ with minor modifications, including a different promotor (chicken β-actin hybrid, CBh) and poly(A) tail. Humanized PC-OxPL-VecTab® sequences were generated to present a lower immunogenic profile and similar or slightly better binding in relation to mouse PC-OxPL-VecTab® formats (unpublished data).

#### AAV generation

AAV5.2 carrying the sequences for the various PC-OxPL-VecTabs® (i.e. the mouse or humanized scFv sequence) or the reporter gene green fluorescent protein (GFP) (i.e. AAV5.2-CBh-GFP) were produced in the Baculovirus-insect cell production system utilizing the artificial intron technology developed by ViroVek Inc ^24^. To this end, two recombinant baculoviruses were generated and subsequently used for AAV production in *Spodoptera frugiperda* Sf9 cells (Invitrogen). The first recombinant Baculovirus (Bac-CapRep) carried both the AAV5.2 Capsid (Cap5.2) and AAV2 Replication (Rep) sequences and was generated using the Bac-to-Bac system (Invitrogen). The second recombinant Baculovirus (Bac-Transgene) carried a transgene cassette containing the mouse, humanized PC-OxPL-VecTab® or GFP transgenes flanked by the AAV2 Inverted Terminal Repeat (ITR) sequences. The baculoviruses with the mE06 and GFP transgenes were produced with the Bac-to-Bac system. In contrast, the baculovirus with the hE06 transgene was produced by homologous recombination in Sf9 cells using the FlashBac- Ultra kit (Oxford Expression Technologies). Recombinant AAV production was performed by co-infection of Sf9 cells with Bac-CapRep and Bac-Transgene. Following harvest, the AAV vectors were purified using two rounds of CsCl density ultra-centrifugation. The CsCl was removed through buffer exchange and the AAV vectors were filter sterilized. Purified AAV vectors were titrated using qPCR.

#### AAV transduction

Human iPSC-derived cultures were transduced with different AAV constructs at a multiplicity of infection (MOI) of 1E05, 1E06 and 1E07 in culturing medium seven to 10 days post-seeding. Half of the medium was removed from the wells before adding the AAV mixture. The same volume of fresh medium was added to the cells at four h post-transduction.

### Gene expression studies

#### RNA extraction and quality control

iCell motor neurons were at a cell density of 350.000-700.000/well using a 12-well plate. At the end of the experiment, lysates were collected in QIAzol lysis reagent (79306, Qiagen). RNA isolation was performed using the miRNeasy Mini kit (217004, Qiagen) according to the manufacturer’s protocol for samples containing <1 μg total RNA.

RNA concentrations were determined using NanoDrop ONE^C^ (Thermo Scientific). Quality control analysis was executed with Bioanalyzer (Agilent RNA 6000 Nano), according to the manufacturer’s instructions. All the samples analyzed contained high-quality RNA, with an RNA integrity number (RIN) above 8.5 and clear ribosomal RNA (rRNA) 18S/28S peaks.

#### DNA extraction

Cell lysates were collected by incubating cells with Accutase and washing themwith D-PBS. DNA isolation was performed using the DNeasy Blood & Tissue kit (69504, Qiagen). DNA concentrations were determined using the NanoDrop ONE^C^ (Thermo Scientific).

#### nCounter Gene Expression (NanoString)

Two pre-designed NanoString Neuroscience panels (Assays and Panels for Neuroscience | NanoString) were selected for gene expression analysis: (a) Neuropathology and (b) Neuroinflammation.

Samples for the nCounter assay were treated as described in the NanoString technologies Gene expression hybridization protocol (Gene Expression CodeSet RNA Hybridization Protocol; MAN- 10056-05) and 5 µL of sample (20 ng/ul) was used in each hybridization reaction. The entire hybridization reaction was used as input for the nCounter prep station and ran on an nCounter MAX cartridge. Each nCounter cartridge was read on the NanoString digital analyzer (nCounter MAX/FLEX) with 280 fields of view (FOV). Finally, digital count files were generated for each sample run.

Raw data was analyzed using the nSolver 4.0 Analysis Software. Raw data files were imported to nSolver 4.0. Datasets were processed according to a two-step method: (1) quality control (QC) and (2) Expression changes.

All samples were verified regarding quality control (QC) parameters using Basic nSolver analysis software (101), applying default settings recommended by NanoString for binding density, mRNA content, mRNA positive normalization factor, mRNA content normalization factor, and fields of view (FOV). Only transcripts with average counts ≥ 80 reads were considered suitable housekeeping genes.

Negative control probe reads were used to define the minimum expression threshold (≥ 15 reads). Therefore, all transcripts with expression levels (Max counts) below the negative control limit were excluded from further analysis. Following QC, samples were clustered into the respective experiment groups. Pairwise ratio comparisons were applied to datasets on both panels to compare experimental groups. Fold-change (FC) was used as a readout for each possible combination. Regression analysis (Coefficient of determination, R^2^) was used to analyze independent replicates, that showed R^2^ ≥ 0.95 across all datasets. Detailed information about the software, including the recommended ranges of values for the different QC parameters, creating experiments, normalization, and analyzing data, can be found at <u>nSolver 4.0 Analysis Software User Manual (NanoString.com).</u>

After processing the datasets, the percentage of transcripts with altered expression following PONPC treatment was calculated based on the initial number of transcripts analyzed. Overlapping transcripts between NanoString panels were signaled, and duplicates were discarded from the final analysis. The percentage of PONPC-sensitive transcripts overlapping with ALS-related genes was evaluated based on the number of ALS-related transcripts in the initial list of NanoString panels. To this end, the number of common transcripts between the ALS datasets and the initial list of NanoString panels was calculated. From these, the number of PONPC-sensitive transcripts was determined.

The effect of AAV5.2-PC-OxPL-VecTab® on transcriptomic changes was assessed on PONPC- sensitive transcripts and transcripts that changed due to transduction (AAV5.2-Control) were excluded from this analysis.

A FC cut-off≥ 10% transcript change was applicable at most analysis changes, except for the assessment of the AAV5.2-PC-OxPL-VecTab® effect, in which any FC towards wt (PSPC) levels was considered therapeutically relevant.

#### Gene ontology (GO) analysis

Gene ontology (GO) analysis, including Fold Enrichment plots, was performed using Shiny GO 0.76^5^. Fold Enrichment was defined as the percentage of genes in the provided gene list that belonged to a certain pathway. The false discovery rate (FDR) was used as a readout on how likely the enrichment found occurred by chance. An FDR cutoff of 0.05 was considered for all GO analyses.

#### cDNA synthesis

Single- and double-stranded DNA was digested with the DNase I, RNase-free kit (EN0525, Thermo Scientific). The reaction was carried out by incubation with DNase I at 37° C for 30 min and stopping the reaction for 10 min at 65° C with EDTA. Reverse transcription was performed with the Maxima First Strand cDNA Synthesis Kit for RT-qPCR (K1641, Thermo Scientific) according to the manufacturer’s instructions. cDNA synthesis was carried out in a 20 µL reaction volume containing 8 uL of RNA, 2 uL of DNase I, and 10 ul of the Maxima mix. The following cDNA synthesis program was used: (25°C, 10 min), (50°C, 15 min), (85°C, five min).

#### Quantitative Reverse Transcriptase Chain Reaction (qRT-PCR)

A PCR master mix was created by adding the appropriate primers and probe. As an endogenous control for PC-OxPL-VecTab® detection, Glyceraldehyde-3-phosphate dehydrogenase (*GAPDH*) (4333764T, Thermo Fisher) was used. The master mix was added to a MicroAmp™ Fast Optical 0.1 mL 96-Well Reaction Plate (4346907, Thermo Scientific). The qPCR reaction was carried out using the QuantStudio™ 5 (Thermo Scientific). The following qPCR program was used: hold (95°C, 20 s), denature (95°C, 1 s), repeated 2-3 (x40) and annealing-elongation (60°C, 20 s).

#### Immunocytochemistry *in vitro* assays

In all immunocytochemistry experiments, cells were initially fixed with 4% (w/v) paraformaldehyde (PFA) (11586711, Thermo Fisher Scientific) for 20 min at RT. Depending on the downstream target, further immunocytochemistry procedures were performed.

#### APOE, pTDP-43, β-tubulin III and mitochondria detection

After washing with PBS (14190-094, Gibco) cells were permeabilized with 0,1% Triton X-100 (M236- 10ML, VWR) for 15min at RT, washed three times with PBS, and incubated with blocking solution (3% BSA [422371X, VWR]) for 30 min at RT. Cells were incubated with the primary antibodies diluted in blocking solution overnight at 4 °C, or for one h at RT. Following three washes in PBS, cells were incubated with the secondary antibody diluted in blocking solution in the dark for 1h at RT. Cells were washed two times with PBS and incubated with Hoechst (H3570, Invitrogen) diluted 1:10000 in PBS in the dark for 10 min at RT. After one wash with PBS, cells were stored at 4 °C until imaging. For phosphorylated TDP-43 (pTDP43) and APOE detection, tris-buffered Saline (TBS) was used instead of PBS. Further antibody details and experimental conditions are listed in detail in **Supplementary Table S2**.

#### PC-OxPL detection

After washing with PBS, cells were permeabilized for 15 min with 0.5% Saponin (47036-50G-F, Sigma), washed three times with PBS, and incubated with blocking solution (5% goat serum [31873, Thermo Fisher Scientific]) for 45 min at RT. Cells were incubated with the humanized E06 full-length IgG1 protein primary antibody (hE06) diluted in blocking solution (1:100) for one h at RT. Following three washes in 1% goat serum, cells were incubated with the goat anti-human IgG Alexa 647 (Invitrogen) diluted in blocking solution (1:750) in the dark for 1 h at RT. Cells were washed two times with PBS and incubated with Hoechst (1:10000 in PBS) in the dark for 10 min at RT. After one wash with PBS, cells were stored at 4 °C until imaging.

### Image acquisition and analysis

#### Imaging

Images of the cells were captured using the ImageXpress Pico system (Molecular Devices) with 10x/20x/40x magnification and up to four detection channels (FITC, TRITC, Cy5, and DAPI), depending on the secondary antibodies and dyes used in each experiment.

#### Analysis

Analyses were carried out using predefined analysis protocols in the CellReporterXpress Software. Average neurite outgrowth was quantified by running the neurite tracing protocol. The percentage of positive cells and all cell average intensities (shown graphically as ‘expression’) were quantified by running the cell scoring protocol. Mitochondria analysis was performed using an ImageJ (v1.53q) plugin to isolate soma-specific mitochondrial signals. This was done by creating a mask (threshold = 32-255) from the original mitochondrial channel and excluding the neurite signal based on size range through the particles’ analysis tool (size 60-100000). The resultant mask, containing the area occupied by mitochondria within the soma, was then merged with the original fluorescence image using the Image Calculator tool, creating an image with mitochondrial fluorescence in the soma region only. This image was analyzed by setting and applying the Measure tool for the Mean Gray value.

#### HTRF TDP-43 Aggregation assay

The HTRF kit (TDP-43 aggregation kit, Cisbio) was used to measure TDP-43 aggregation, which is defined as insoluble cytoplasmic inclusions, according to the manufacturer’s instructions. Briefly, 125.000/25.000 iCell Motor Neurons/Astrocytes (Fujifilm) were seeded in a co-culture setting on a Cytoview 96-well plate (M768-tMEA-96B, Axion Biosystems). Cells were cultured in BrainPhys^TM^ Neuronal Medium (05790, STEMCELL TECHNOLOGIES), supplemented with and without DAPT (week one and two-three, respectively). AAV transduction was performed on day seven post-cell seeding, and PSPC or PC-OxPL treatment (25 µM) was performed on day 21 for 24 h, followed by 24 h washout. On days 21-22 *in vitro*, two cellular lysates (wells) were pulled for each condition, and the assay ran with three technical replicates. For each sample, the ratio of acceptor and donor emission signals was calculated as follows: Ratio = (Signal 665 nm/ Signal 620 nm) x 10E4. The aggregated ratio was calculated for each sample according to the following: Aggregated Ratio = Ratio Disaggregated Sample/Ratio Control Sample.

## Animal Experiments

### sALS CSF mouse model

Experiments were performed as described in Wong et al. ^26^, with minor modifications.

#### Patient demographics and CSF collection

CSF was obtained from one sALS patient diagnosed by board-certified neurologists with a subspeciality interest in ALS. CSF from one sALS patient seen seen at the Larry G. Gluck Division of ALS Research at the Tisch MS Research Center of New York (Tisch Center) was selected for this study. Information on patient demographics can be found in ^26^.

Institutional Review Board approval and informed consent according to the Declaration of Helsinki was granted prior to CSF collection. Samples were collected using sterile techniques either by lumbar puncture or access port aspiration of surgically implanted pumps. CSF samples were centrifuged at 200 x g for 15 min to remove cells, confirmed to be free of red blood cell contamination by microscopy, and then stored in aliquots at –80°C.

#### Intrathecal injection into cervical subarachnoid space

Adult female C57BL/6J mice (aged nine weeks at the time of first surgery) purchased from The Jackson Laboratory (Bar Harbor, ME) were used. All procedures were approved by the Institutional Animal Care and Use Committee at Mispro Biotech Services (New York). Prior to surgery, mice were anaesthetized with ketamine (110 mg/kg) and xylazine (10 mg/kg) cocktail and received subcutaneous injections of 0.1 mg/kg buprenorphine, 2.5 mg/kg baytril and 1 mL 0.9% saline. Laminectomies at cervical levels 4 (C4) and 5 (C5) were performed to expose the underlying spinal cord. A 32-gauge Hamilton syringe was inserted underneath the dura mater and 5 µL of AAV5.2-PC-OxPL-VecTab® (1.3E12 gc/mouse) or saline were slowly injected into the subarachnoid space. After four weeks, a second surgery was performed and 3 µL of sALS CSF or saline were injected into the subarachnoid space. At least three mice were injected per group at each round and three independent rounds were performed. Mice were assigned to different treatment groups in a randomized manner.

#### Preincubation of PC-OxPL-VecTab® with sALS CSF prior to intrathecal injection

To allow preincubation of sALS CSF, Control-VecTab® or PC-OxPL-VecTab® were produced at a concentration > 0.5 mg/mL. All buffers used were made using Versylene (endotoxin-free and sterile) water. Endotoxin was removed from instruments by incubation with 0.1 M NaOH for at least 16 h. HEK293E-253 cells were transfected with endotoxin-free plasmid DNA containing the sequences of interest using the rPEx technology (RPEXBIO). At six days post transfection conditioned medium containing recombinant protein was harvested by centrifugation and the samples stored at 4 °C. Using IMAC purification, the recombinant proteins were bound in batch to 0.5 mL Nickel Excel Sepharose for four to five h at 20 °C. Then, Nickel Excel Sepharose containing bound protein was harvested by centrifugation and transferred into a gravity flow column. A-specific bound proteins were removed by washing the column with IMAC buffer A containing 0 mM and 10 mM imidazole. The protein was eluted with IMAC buffer A containing 500 mM imidazole and collected into 2.5 mL fractions. Recombinant protein-containing fractions were pooled. The conditioned medium and the unbound IMAC fraction were analyzed by LabChip capillary electrophoresis. Next, the buffer was exchanged for PBS by desalting using a HiPrep 26/10 desalting column equilibrated in PBS. Recombinant protein-containing fractions were pooled. If necessary, the sample was concentrated using an Amicon4 10 kDa spin filter. For formulation purposes, the pool was sterilized by filtration over a 0.22 μm syringe filter, and the product was stored in 0.5 mL aliquots at 4°C. Finally, both scFvs were analyzed by Labchip capillary electrophoresis.

Adult female C57BL/6J mice (aged 13 weeks at the time of surgery) purchased from The Jackson Laboratory (Bar Harbor, ME) were used. Prior to intrathecal injection, two µg of purified recombinant scFv control-VecTab® or PC-OxPL-VecTab® were mixed with either saline or sALS CSF and kept on ice then at RT for 10 min prior to injection. The intrathecal procedures were performed as described above. Two to five mice were injected per experimental group and one round of experiments was performed.

### Behavioral testing

#### Motor deficit score testing

Following intrathecal delivery of CSF, all mice underwent motor testing at one day post-injection (DPI). Forelimb reaching, gripping and tail flaccidity were evaluated on a three-point scale. Mice were held by their tails above their cage bars and allowed to reach and grip the bars for five trials. Mice displaying no motor deficits were given a score of 0. Any deficits in either reaching or gripping were each given a score of 1. Specifically, inaccurate reach was considered a reaching deficit, and weakness in grip strength or clenched forepaws were scored as gripping deficits. Tail flaccidity was also given a score of 1. All motor testing was performed blinded with respect to treatment groups.

#### Grip strength testing

Mice were habituated to the grip strength meter (TSE systems) for three days prior to surgery. Each mouse was given one min to explore the grip strength meter, then held by their tails and allowed to grip the bar with both forelimbs for five consecutive trials. After a 30 s rest period, the mice were given another five trials to grip and then returned to their home cage. Baseline grip strength force was measured at one day prior to surgery and grip strength was also measured at one DPI. The mean grip strength force was calculated from five trials. Normalized grip strength values were calculated by dividing mean grip strength force on post-injection testing day by mean baseline grip strength force.

#### Tissue harvesting

Mice were overdosed with ketamine (300 mg/kg) and xylazine (30 mg/kg) and then perfused transcardially with phosphate-buffered saline (PBS) followed by 4% paraformaldehyde in 0.1 M PBS, pH 7.4. Spinal cords were dissected, postfixed in 4% paraformaldehyde overnight and then placed in 30% sucrose overnight for cryoprotection.

Cervical spinal cords were cut half cm rostral and half cm caudal to the injection site. The one cm segments were then embedded and frozen in Tissue Tek® (VWR International, PA). Spinal cords were sectioned sagittally at 20 µm thickness using a cryostat (Leica) and then slide-mounted onto Histobond® slides (VWR International, PA). The anatomical orientation of tissue sections and the order and position in which they were mounted onto the slides were kept consistent to facilitate unbiased histological comparisons, as described in further detail below.

#### Immunohistochemistry sALS CSF mouse model

Immunostaining was performed on a series of spinal cord sections at 100 µm intervals throughout the cervical spinal cord. Details on the antibodies used and experimental conditions are listed in **Supplementary Table S2**.

#### ChAT, TDP-43, GFAP and IBA1 detection

Slides with spinal cord sections were washed three times in 0.1% triton X-100 in PBS (PBS/T), then incubated in 10% normal goat serum (NGS) or normal donkey serum (NDS) in PBS/T for 1 h at RT. Primary antibodies were diluted in 10% NGS or NDS in PBS/T and incubation occurred overnight at 4°C. After incubation, slides were rinsed three times in PBS and incubated in the appropriate Alexa- Fluor secondary antibodies (Invitrogen) in 10% NGS or NDS in PBS/T for 1.5 h at RT. Slides were rinsed three times in PBS and then counterstained with 1:2500 DAPI in PBS (Invitrogen) for five min. After two final washes in PBS, free-floating brain sections were mounted onto slides, and slides were mounted using Fluoromount (Sigma).

#### PC-OxPL detection

Slides with spinal cord sections were washed three times in PBS, then incubated in blocking buffer (10% NGS in PBS/T) for one h at RT. The hE06 protein was used to detect PC-OxPL, which was diluted blocking buffer and incubated overnight at 4°C. After incubation, slides were rinsed three times in PBS for 10 min and incubated in the appropriate dilution of goat anti-human IgG Alexa 647 (Invitrogen) in blocking buffer for 1.5 h at RT. Slides were rinsed three times in PBS and then counterstained with 1:2500 DAPI in PBS (Invitrogen) for five min. After two final washes in PBS, slides were mounted using Fluoromount (Sigma).

#### Histological analyses

Images were captured at 20X magnification using a Zeiss Axio Imager. Acquisition parameters and exposure times were kept consistent for each antibody stain. To ensure unbiased comparisons between experimental groups, spinal cord images were captured from similar tissue section numbers on the slides and matching anatomical regions were verified by experimenters. The number of motor neurons and immunostaining intensities were quantified using ImageJ software. Three images were quantified per mouse for cell counts to calculate the mean motor neuron number. Fluorescence intensities were measured as mean grey values in regions of interest. Both imaging and quantification were performed by experimenters blinded to treatment groups.

### Wt Mini pigs

#### Intrathecal delivery of AAV and tissue collection

All experiments were carried out according to the guidelines for the care and use of experimental animals and approved by the State Veterinary. Ten nine-month-old wt mini pigs were selected at the Institute of Animal Physiology and Genetics in Libechov (Czech Republic). For intrathecal administration, under deep sedation and after cleaning and disinfection of skin in the lumbar region of animals, a polyethylene tubing (PE10, 427401, Intramedic Clay Adams) attached to a 5-mL syringe passed through a spinal needle (Spinocan; 4501195-13; 1.1 88 mm; B. Braun Melsungen) was used. After tubing insertion into the intrathecal space, approximately one to two mL of CSF was collected to ensure precise localization of the tubing. Next, four mL of AAV5.2-CBh-GFP (4.5E13 gc/mL) or AAV5.2-CBh-PC-OxPL-VecTab® (2.9E13 gc/mL) were injected in five mini pigs per group. After administration all animals in all groups were under observation until awakening. For at least three days post-application, animals were delivered daily analgesics for pain relief.

Eight weeks after treatment, mini pigs were euthanized under deep sedation followed by anesthesia overdosing with Propofol and whole-body PBS perfusion of the animal. Once removed from the skull, the brain was coronally sliced into three to four mm blocks from frontal to caudal. The right hemisphere was used for biomolecular analyses, where 38 brain punches were taken, snap-frozen in liquid nitrogen and stored at 80°C. The right hemisphere was fixed by 4% paraformaldehyde (PFA) overnight and then transferred to 30% sucrose with sodium azide for sectioning. The spinal cord was separated into the cervical, thoracic and lumbar segments and each segment was divided into three pieces, two were snap-frozen in liquid nitrogen for molecular analysis, and one was PFA-fixed and transferred to 30% sucrose with sodium azide for immunohistochemistry purposes.

#### DNA isolation and vector (vDNA) quantification

DNA isolation from brain tissue punches was performed using the DNeasy blood and tissue kit (QIAGEN, Germany). DNA concentrations were determined using the NanoDrop ONEc (Thermo Scientific). Primers specific for the CBh promoter sequence were used to measure the vector genome copies (gcs) by TaqMan qPCR (Thermo Fisher Scientific). The amount of vector DNA was calculated based on a plasmid standard curve. Results were reported as gc/ug genomic DNA.

#### RNA isolation and transgene mRNA quantification

RNA isolation was performed using the RNeasy Mini kit (74104, Qiagen) according to the manufacturer’s protocol and eluted in 30 µl of RNase-free water. RNA concentrations were determined using NanoDrop ONEC (Thermo Scientific).

cDNA synthesis and qPCR were performed using the same protocol as for the cells (see above). Real-time PCR amplification was performed with customized TaqMan primers for GFP or PC-OxPL- VecTab® expression and with TaqMan primers for Sus scrofa Hypoxanthine Phosphoribosyltransferase 1 (*Hprt1*) porcine housekeeping gene (Ss03388274_m1, Thermo Fisher). The transgene mRNA expression levels were calculated as FC to ss *Hprt1* expression.

#### Immunohistochemistry wt mini pig study

PFA-fixed brain slices were cut into 20-mm sections (10–15 sections/slice), spread on a microscopic slide and air-dried. For immunostaining analysis, the sections on glass were antigen-retrieved by treating in citrate buffer (pH = 6) at 90°C for 30 min. Subsequently, endogenous peroxidase activity was blocked with a solution of 0.3% hydrogen peroxide in methanol for 20 min. The brain sections were immunostained using the rabbit primary antibody anti-GFP (1:1.000), incubated with a biotinylated donkey anti-rabbit secondary antibody (1:400, RPN 1004V; GE Healthcare), followed by an avidin-peroxidase complex (1:400, A3151; Sigma-Aldrich). The avidin-peroxidase complex was visualized by incubation with a solution containing a dissolved 3,30-diaminobenzidine tablet (4170; Kementec Diagnostics). The sections were dehydrated and mounted with DePeX (Sigma). Images were acquired using a histological scanner (VS120-5 Virtual Slide Microscope fluorescence; Olympus).

## Statistical analysis

Statistical analysis was performed with GraphPad Prism (versions 9.2.0 and 10.2.3).

Differences in (Ox)PCs measured by LC-MS in CSF and plasma were determined using a two-tailed Welch’s t-test.

Differences in neurite outgrowth (TUBB3), PC-OxPL and pTDP-43 profile as measured by immunohistochemistry were evaluated using two-way ANOVA. Differences in the TDP-43 aggregation profile were assessed using one-way ANOVA with Tukey’s or Dunnett’s multiple comparison test.

Differences in PC-OxPL content on different apolipoproteins were assessed with a one-way ANOVA with Dunnett’s multiple comparison test. Differences in the APOE profile as measured by immunohistochemistry were evaluated using a two-way ANOVA with Šidák’s multiple comparison test. Normalized APOE concentrations in motor neurons, plasma, and CSF were determined according to a two-tailed Mann-Whitney test.

Motor deficit scores, normalized grip strength, motor neuron numbers, and immunostaining intensities were analyzed using a one-way ANOVA with Bonferroni post hoc analyses.

All values are expressed as mean ± the standard error of the mean (SEM). No statistical predictions were used to determine sample size (n) but the n used resembled published literature and what is generally performed in the field. Statistical significance was considered for p-values (p) ≤ 0.05 (p < 0.0001: ****; p < 0.001: ***; p < 0.01: **; p < 0.05: *; p ≥ 0.05: ns or not significant).

## Results

### PC-OxPL accumulates in the diseased ALS brain and spinal cord

TDP-43 pathology is present in 97% of all ALS patients and spreads differentially throughout brain areas with disease progression ^27^. We assessed the presence of PC-OxPL across different human brain regions in patients with ALS. Evaluation of spinal cord, frontal cortex, thalamus, hippocampus and occipital cortex areas showed the presence of PC-OxPL, primarily within cell bodies and with variable signal intensities, depending on the brain area and individual (**Fig. 1a**). Overall, more PC-OxPL signal was retrieved in the spinal cord ventral horn and white matter areas of ALS patients in comparison to non-demented controls (NDC) (**Fig. 1b**). We then investigated the subtype cell specificity of PC-OxPL accumulation by analyzing the hE06, ChAT (motor neuron) and DAPI (all cells) signal profile in the spinal cord. We found specific PC-OxPL accumulation in ChAT+ neurons in the spinal cord ventral horn regions, as shown by the co-localization of hE06 with ChAT, but not with DAPI (**Fig. 1c-d**). Next, we characterized the profile of PC-OxPL species in ALS. To address this, targeted lipidomics analysis was performed on CSF and plasma collected from ALS patients and HC. We focused on six precursor families (PC) and their corresponding PC-OxPL products. Analysis of CSF collected from ALS patients revealed an overall increase in 16/18 PC-OxPL detected species, out of which 10/18 found to be significantly changed (**Fig. 1e**). Notably, this upregulation trend was replicated in their precursors profile in comparison to HC (**Fig. 1e**). The same species displayed an opposite profile in matched ALS plasma, with 12/18 found to be significantly downregulated in comparison to HC (**Fig. 1f**).

**Figure 1.**
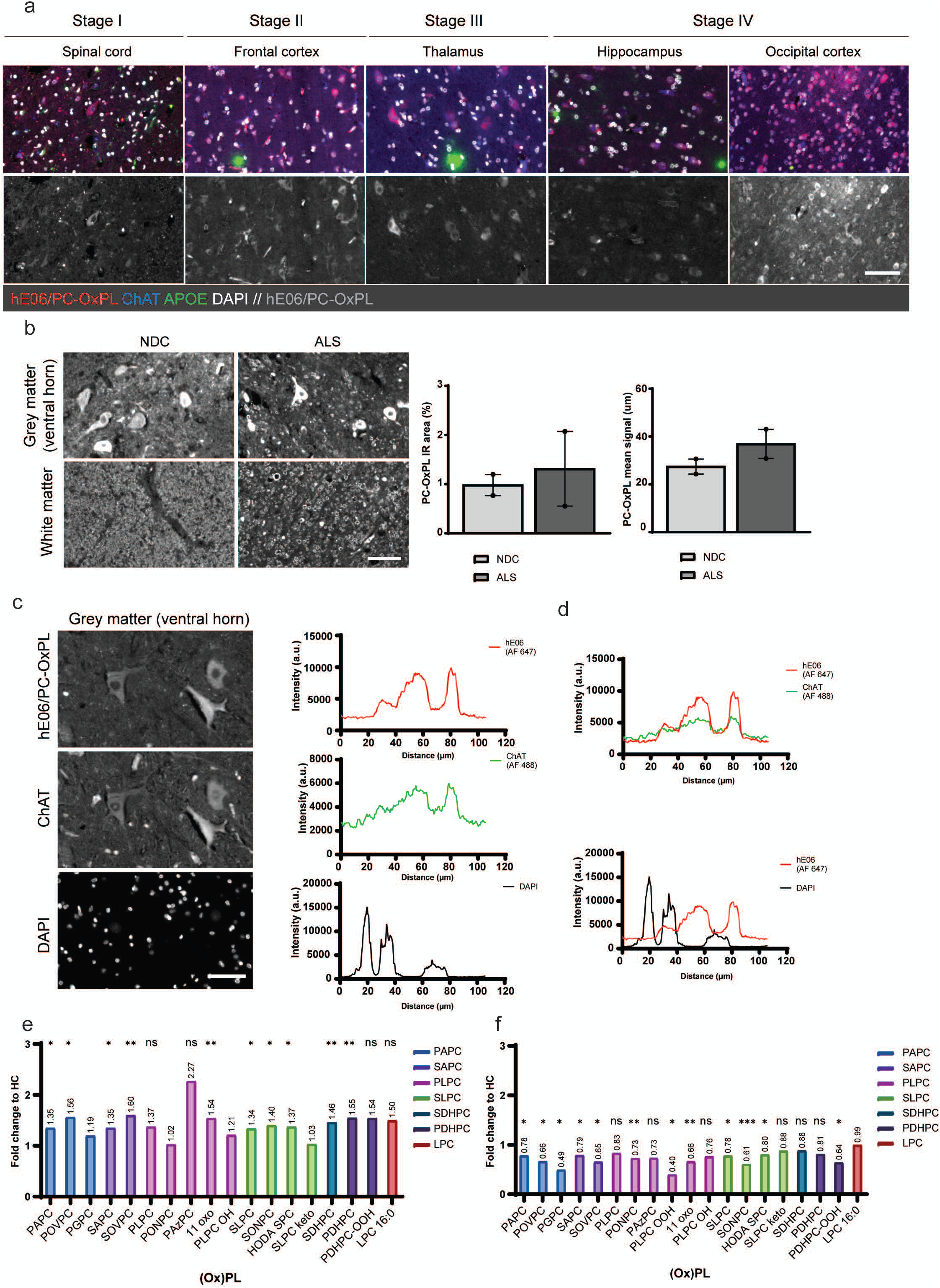
PC-OxPL accumulates in the brain and spinal cord of ALS patients. **a.** Representative images of PC-OxPL pathology, as recognized using the hE06 full-length antibody, across the different stages that characterize ALS pathology and progression ^27^. Scale bar = 100 µM. **b.** *Left panel.* Representative images of PC-OxPL accumulation in the spinal cord of ALS patients and non-demented controls (NDC). Scale bar = 100 µM. *Right panel.* Quantification of PC-OxPL accumulation in the spinal cord of ALS patients and non-demented controls (NDC), as per infrared (IR) area and mean signal intensity. Values are expressed as means ± SEM. N = 2 individuals per group. **c.** *Left panel.* Representative images of PC-OxPL and ChAT co-localization in the grey matter region of the spinal cord. Scale bar = 100 µM. *Right panel.* Line profiling of hE06/PC-OxPL and ChAT or DAPI co- localization in the grey matter region of the spinal cord. The profile is plotted as the intensity of each channel (AF 647, AF 488 or DAPI) per pixel. **d.** Co-localization profile of hE06/PC-OxPL with ChAT (*upper panel*) and DAPI (*lower panel*). **e.** FC of individual (Ox)PC species in CSF from ALS patients compared to HC. Values are expressed as means (n = 15). **f.** FC of individual (Ox)PC species in plasma from ALS patients compared to HC. Values are expressed as means (n = 15). Individual data points/SDs are omitted in **e.** and **f.** for clarity. Two-tailed Welch’s t-test, ***p < 0.001: **p < 0.01, *p < 0.05, ns, p ≥ 0.05.

### PC-OxPL induces ALS-like phenotypes in healthy motor neurons

Having established the pathological PC-OxPL phenotype in ALS, we aimed to understand what molecular mechanisms are activated by PC-OxPL in relevant brain cell types. 16:0/18:1, *1-palmitoyl- 2-oleoyl-sn-glycero-3-phosphocholine* (POPC), is found to be the most abundant PC species in the rat, porcine and human brain ^28–30^ and its oxidation results in two main 16:0/09:0 PC-OxPL species: PONPC and PAzPC. Hence, we compared the effects of PONPC and PAzPC on wt motor neurons. Both PC- OxPL species showed clear effects on neuronal toxicity compared to a non-oxidized PC (PSPC). Yet PAzPC showed a higher level of toxicity leading to irreparable cell death which could compromise further analysis (**Supplementary Fig. S1a-b**). Therefore, we proceeded with PONPC as the selected PC-OxPL agent for subsequent *in vitro* assays.

Transcriptomic analysis of iPSC-derived motor neurons using NanoString Technology was performed to understand the molecular mechanisms activated by PC-OxPL, using two available panels: ‘Neuropathology’ and ‘Neuroinflammation’. Of the 770 transcripts, exposure of wt neurons to PONPC led to 173 (Neuropathology) and 138 (Neuroinflammation) differently expressed (DE) genes, accounting for approximately 25% transcriptome alterations in response to PONPC exposure (**Supplementary Table S4**). Gene Ontology (GO) analysis of the same gene list revealed ‘Response to stimulus’, ‘Programmed cell death’, ‘apoptosis’, ‘regulation of molecular functions’, ‘transcriptome remodeling’ and ‘protein phosphorylation’ to be among the most representative pathways in motor neurons following PONPC exposure (**Fig. 2a**). Next, we used commercially available iPSC-derived motor neurons harboring known ALS-associated mutations to replicate part of ALS pathophysiology *in vitro* ^31^. The transcriptomic profile of extensively characterized ALS motor neurons carrying TDP- 43^M337V^ and SOD1^G93A^ was compared to one of wt motor neurons exposed to PONPC. Overlapping dysregulation in gene expression was observed between TDP-43^M337V^ and SOD1^G93A^ transcriptome (NP = 35.08%; NI = 34.35%), as was expected considering that these genes are related to ALS pathogenesis (**Fig. 2b, Supplementary Table S4**). Considerable transcriptome overlap was found between wt motor neurons exposed to PONPC and SOD1^G93A^ motor neurons (NP = 39.69%; NI = 50.31%) and, to a lesser extent, TDP-43^M337V^ motor neurons (NP = 28.48%; NI = 50.31%) (**Fig. 2b, Supplementary Table S4**). Of all DE transcripts, 31.03% (NP) and 22.58% (NI) were found to be deregulated among all three iPSC-derived motor neuron lines (wt exposed to PONPC, TDP-43^M337V^ and SOD1^G93A^). To further validate these findings in a broader ALS context and to exclude effects of general oxidative stress and inflammatory hallmarks ^32^, we compared the changes observed in wt motor neurons exposed to PONPC with two independent datasets: (1) ALSoD ^33^ and (2) Postmortem spinal cord tissue from patients with sporadic ALS (D’Erchia et al. ^34^ (**Supplementary Table S5**) (**Fig. 2c**). Our analyses revealed that 5% (n = 36) of the transcripts that were previously related to ALS according to selected databases (1, 2) clearly altered expression following PONPC treatment ^33,34^ (**Fig. 2d**). This is likely an underestimation as the NanoString panels include a limited number of genes (n = 770), instead of a complete transcriptome analysis. The overlapping transcripts between databases (1, 2) and with altered expression following PONPC exposure in motor neurons are listed in **Table 1**, along with a brief description of their function.

**Figure 2.**
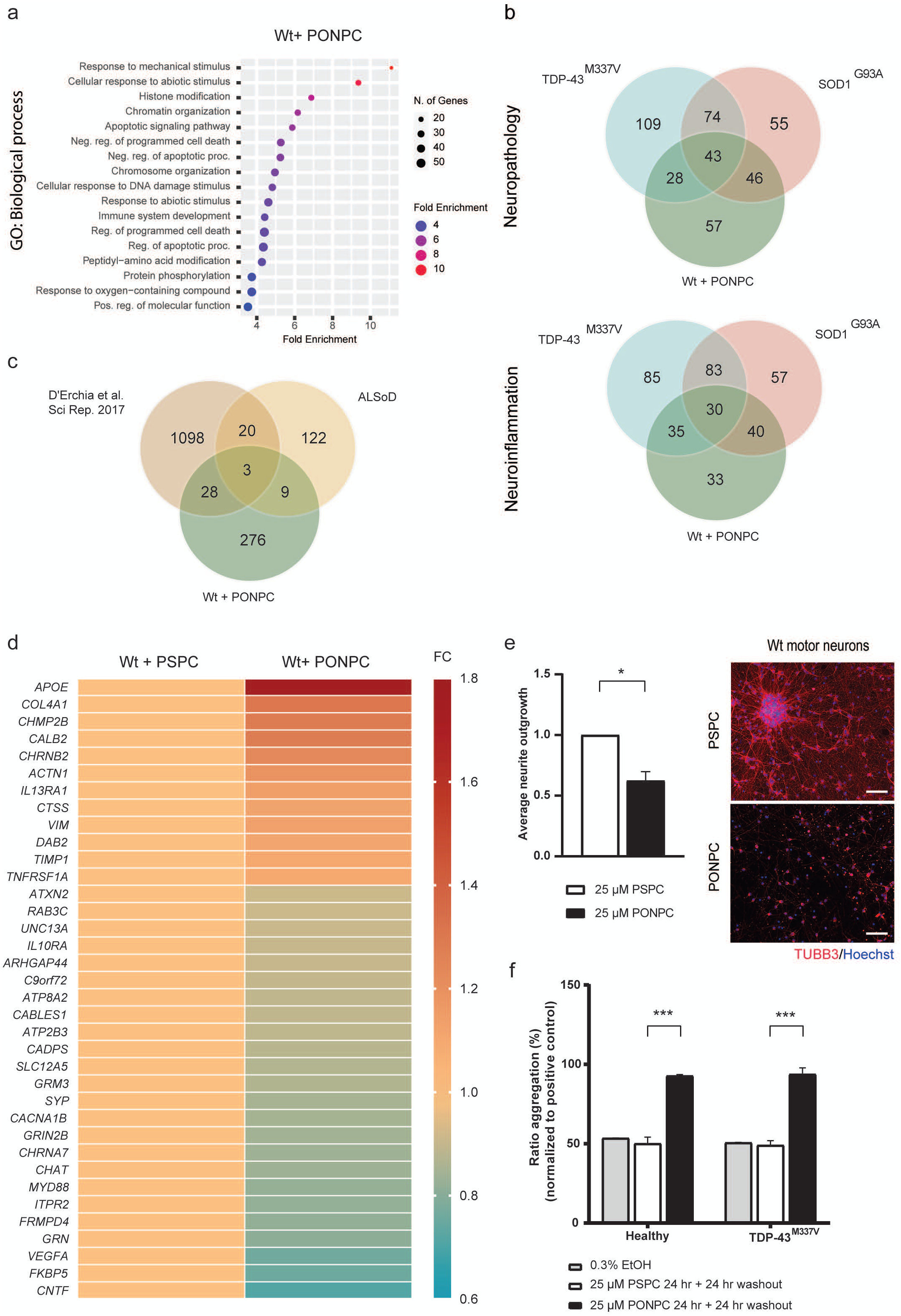
PONPC induces ALS-like signatures in motor neurons. **a.** Dot plot representation of the 20 most enriched independent processes among the DE genes in wt motor neurons exposed to PONPC. Gene expression data from both Neuroinflammation and Neuropathology were used in the analysis. All pathways are sorted by Fold Enrichment value. **b.** Venn diagram depicting overlapping transcripts changed following PONPC treatment of wt and ALS (TDP-43^M337V^ and SOD1^G93A^) motor neurons according to the deregulated neuropathology (*top*) and neuroinflammation (*bottom*) NanoString nCounter panels. **c.** Venn diagram showing ALS-like transcriptome signatures in wt neurons exposed to PONPC. D’ Erchia et al (2017) ^34^ and ALSoD ^33^ were used as reference ALS-relevant databases for overlap comparison. **d.** Heatmap representation of PONPC/ALS-related expression changes following wt motor neuron exposure to PSPC or PONPC. Transcripts derived from neuropathology NanoString nCounter panels overlap with ALS databases. Expression changes are displayed as FC and normalized to the wt + PSPC condition. **e.** PC-OxPL (PONPC) treatment of wt motor neurons impairs neuronal outgrowth. *Left panel:* Values are expressed as means ± SEM and were normalized to 25 µM PSPC condition. Two-way ANOVA, *p=0.0368; n = three independent experiments, one to four replicates. *Right panel:* Representative images showing decreased neuronal outgrowth following PC-OxPL (PONPC) treatment of Axol motor neurons. Scale bar = 100 µM (Tuj-1 (Neuron-specific class III beta- tubulin), Cy5). **f.** TDP-43 aggregation in wt and TDP-43^M337V^ motor neurons exposed to PSPC or PC- OxPL. TDP-43 aggregation is shown as the ratio of aggregation measured by the HTRF TDP-43 assay. Values are expressed as means ± SEM, normalized to the assay positive control. Ordinary one-way ANOVA with Tukey’s multiple comparison test, ***p < 0.001; n= two independent experiments, six replicates.

**Table 1:** PONPC-sensitive transcripts in motor neurons with a role in ALS. <u>The Human Protein</u> <u>Atlas</u> ^35^ was used to describe gene names and functions.

| Gene | Description | Function |
| --- | --- | --- |
| *ACTN1* | Actinin alpha 1 | Actin binding |
| *APOE* | Apolipoprotein E | Lipid metabolism, cholesterol metabolism, lipid transport, steroid metabolism |
| *ARHGAP44* | Rho GTPase activating protein 44 | GTPase activation, exploratory dendritic filopodia |
| *ATP2B3* | ATPase plasma membrane Ca<sup>2+</sup> transporting 3 | Intracellular calcium homeostasis |
| *ATP8A2* | ATPase phospholipid transporting 8A2 | Lipid flipping, lipid transport |
| *ATXN2* | Ataxin 2 | Endocytosis, mTOR signals, ribosomal translation, mitochondrial function |
| *C9orf72* | C9orf72-SMCR8 complex subunit | Autophagy, endosomal trafficking, endocytic transport |
| *CABLES1* | Cdk5 and Abl enzyme substrate 1 | Neuronal outgrowth |
| *CACNA1B* | Calcium voltage-gated channel subunit alpha1 B | Calcium neuronal signaling |
| *CADPS* | Calcium dependent secretion activator | Vesicular exocytosis of neurotransmitters and neuropeptides |
| *CALB2* | Calbindin 2 | Calcium signaling |
| *CHAT* | Choline O-acetyltransferase | Biosynthesis of acetylcholine |
| *CHMP2B* | Charged multivesicular body protein 2B | Endosomal transport, lysosomal degradation of membrane proteins, lysosomal enzymes, and lipids |
| *CHNA7* | Cholinergic receptor nicotinic alpha 7 subunit | Acetylcholine receptor, ligand-gated ion channel |
| *CHNB2* | Cholinergic receptor nicotinic beta 2 subunit | Acetylcholine receptor, ligand-gated ion channel |
| *CNTF* | Ciliary neurotrophic factor | Survival factor, prevention of motor axons degeneration |
| *COL4A1* | Collagen type IV alpha 1 chain | Angiogenesis, extracellular matrix homeostasis |
| *CTSS* | Cathepsin S | Protein degradation |
| *DAB2* | DAB adaptor protein 2 | Apoptosis, Differentiation, Endocytosis, Protein transport, Wnt signaling pathway |
| *FKBP5* | FKBP prolyl isomerase 5 | Immunoregulation, protein folding and trafficking |
| *FRMPD4* | FERM and PDZ domain containing 4 | Excitatory synaptic transmission |
| *GRIN2B* | Glutamate ionotropic receptor NMDA type subunit 2B | Calcium permeability, synaptic plasticity |
| *GRM3* | Glutamate metabotropic receptor 3 | G-protein coupled receptor for glutamate |
| *GRN* | Granulin precursor | Lysosomal regulation, Inflammatory response |
| *IL10RA* | Interleukin 10 receptor subunit alpha | Inflammatory response |
| *IL13RA1* | Interleukin 13 receptor subunit alpha 1 | Inflammatory response |
| *ITPR2* | Inositol 1,4,5-trisphosphate receptor type 2 | Calcium transport |
| *MYD88* | MYD88 innate immune signal transduction adaptor | Inflammatory response |
| *RAB3C* | RAB3C, member RAS oncogene family | Protein transport |
| *SLC12A5* | Solute carrier family 12 member 5 | Potassium-chloride cotransport |
| *SYP* | Synaptophysin | Synaptic plasticity |
| *TIMP1* | TIMP metalloproteinase inhibitor 1 | Metalloproteinase inhibitor, cell proliferation, anti-apoptotic function |
| *TNFRSF1A* | TNF receptor superfamily member 1A | Inflammation, apoptosis |
| *UNC13A* | Unc-13 homolog A | Neurotransmitter release |
| *VIM* | Vimentin | Cell shape maintenance, cytoskeletal homeostasis, cytoplasm integrity |

Given the increasing focus on TDP-43 loss-of-function in ALS ^36^ we investigated the overlap in signatures between PONPC-sensitive transcripts and the genome landscape following neuronal TDP- 43 depletion. We used an available RNA-sequencing dataset of iPSC-derived neurons displaying reduced TDP-43 expression following CRISPRi approach ^37^. Of the total PONPC-sensitive transcripts, approximately 40% (111/311) were found to be altered in neurons with reduced TDP-43 expression (**Supplementary Table S5**). Among other relevant genes, we found *UNC13A*, known to underline ALS pathology ^38^ and further strengthen the relevance of PC-OxPL neurotoxicity in ALS.

Because ferroptosis is closely linked to lipid oxidation ^39^, we investigated the presence of ferroptosis signatures in the transcriptomic data. We cross-checked both the (1) DE transcripts following wt motor neurons + PONPC and (2) ALS-related DE transcripts following wt motor neurons + PONPC with FerrDb, an online curated database for ferroptosis markers and associations ^40^. Significant ferroptosis-related hits were found among the DE following wt motor neurons + PONPC (**Supplementary Table S6**) while no statistically significant ferroptosis hits were found among the ALS-related DE following wt motor neurons + PONPC (**Supplementary Table S6**).

Next, we addressed the phenotypic consequences of motor neuron exposure to PONPC, by focusing on general neuronal health, TDP-43 aggregation and phosphorylation ^41,42^. Exposure of wt motor neurons to PONPC led to neuronal toxicity, as shown by changes in neurite outgrowth and network organization (**Fig. 2e**). Notably, PONPC exposure caused marked and clear TDP-43 aggregation both in wt and ALS (TDP-43^M337V^) motor neurons after 24 h of PONPC exposure followed by a 24 h washout (**Fig. 2f**). In general, the effects were comparable for both iPSC-derived motor neuron lines and more significant than the effect of the commonly used proteasome inhibitor carbobenzoxy-Leu-Leu-leucinal (MG-132) (**Supplementary Fig. S2a**). No significant alterations in TDP-43 phosphorylation were observed under similar conditions (**Supplementary Fig. S2b**). Finally, mitochondrial dysfunction is an essential marker of oxidative stress and, particularly, of cellular ferroptosis ^39^. Yet, we found no differences in the mitochondrial phenotypes of motor neurons following PONPC exposure according to selected readouts (**Supplementary Fig. S2c-d**).

In summary, PONPC-exposed motor neurons developed ALS-associated transcriptomic changes, extensive TDP-43 aggregation, and altered neurite outgrowth, all of which are established ALS hallmarks.

### APOE is the main carrier of PC-OxPL in the CNS

*APOE* was among the most DE genes in wt and ALS motor neurons in response to PONPC exposure (**Table 2**; **Fig. 3a**). This pattern was mirrored by increased protein expression, as shown by the increase in the number of APOE+ cells in a time-dependent manner (**Fig. 3b, left panel**) and average intensity (**Fig. 3b, right panel**). To better understand the apolipoprotein/PC-OxPL profile in ALS we analyzed biofluids from ALS patients and the respective PC-OxPL content across different apolipoproteins (APOB, APO(A), APOE, APOC-III, APOA-I). PC-OxPL were detected predominantly on APOE particles in the CSF derived from ALS patients (**Fig. 3c**), and this was replicated in HC biofluids (**Supplementary Fig. S2e**). A distinct profile was observed in the plasma collected from the same patients, where a homogenous distribution of PC-OxPL on the different apolipoproteins was observed, with the exception of APOA-I (**Fig. 3c**). We further investigated the APOE pattern and profile in ALS. Quantitative and qualitative analysis revealed increased APOE expression in white and grey matter regions of the ALS spinal cord in comparison to NDC (**Fig. 3d**). Similarly to the trend observed in the apolipoprotein/PC-OxPL profile, APOE concentration was significantly increased in CSF, but not plasma, of ALS patients in comparison to HC (**Fig. 3e**). Taken together, these results place APOE at the center of PC-OxPL metabolism in the CNS.

**Figure 3.**
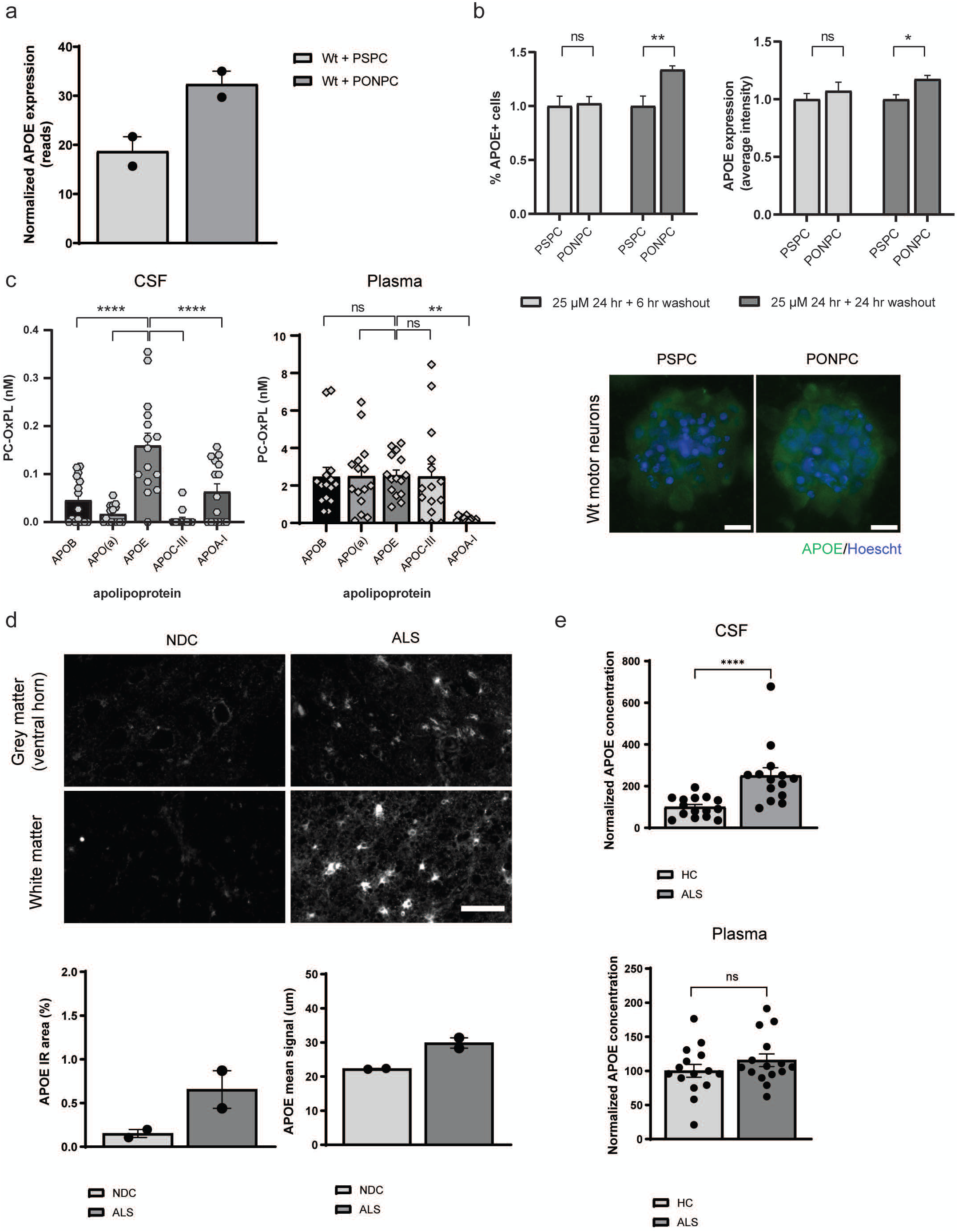
Exposure to PC-OxPL triggers *APOE* expression, which is the main carrier of PC-OxPL in the CSF. **a.** *APOE* transcript expression in motor neurons exposed to either PSPC or PONPC, as determined by NanoString nCounter analysis. Values are expressed as means ± SEM. **b.** *Upper panel:* Elevated APOE profile (% positive cells and average intensity) in wt motor neurons exposed to PONPC according to different washout periods. Values are expressed as means ± SEM. Two-way ANOVA with Šidák’s multiple comparisons test, **p=0.0064, *p = 0.0167, ns, p ≥ 0.05; n = four, two independent experiments. *Lower panel:* Representative images showing increased APOE profile in wt motor neurons at 24 h PONPC treatment + 24 h washout. Scale bar = 20 µM (APOE, FITC). **c.** PC-OxPL levels as detected on the surface of different apolipoprotein particles as measured in paired CSF and plasma from ALS patients. Values are expressed as means ± SEM and each dot represents a single individual (n = 15). One-way ANOVA with Dunnett’s multiple comparisons test, ****p < 0.0001, ***p = 0.0007, **p = 0.0015, ns, p ≥ 0.05. **d.** *Upper panel.* Representative images of APOE profile in the spinal cord regions (grey and white matter) of NDC and ALS individuals. Scale bar = 100 µM. *Lower panel.* APOE expression, as determined by infrared (IR) area and mean signal intensity in the spinal cord regions (grey and white matter) of NDC and ALS individuals. Values are expressed as means ± SEM; n = NDC or ALS individuals per group. **e.** APOE concentrations as detected in paired CSF and plasma from ALS patients compared to HC. Values are expressed as means ± SEM and each dot represents a single individual (n = 15). Mann-Whitney U test, ****p < 0.0001, ns, p ≥ 0.05.

### AAV5.2-delivered PC-OxPL-VecTab®, prevents neurotoxicity in wt motor neurons

The characterization assays suggest that PC-OxPL neutralization is a promising approach to target ALS-related phenotypes. The following sections describe the development, characterization, and viability of a vectorized antibody treatment to target PC-OxPL neurotoxicity.

We developed an AAV5.2-delivered approach based on a vectorized scFv, binding to the PC headgroup of oxidized phospholipids (PC-OxPL-VecTab®) (**Fig. 4a**) ^48^. AAV5.2-GFP successfully transduced ALS-relevant cell types including motor neurons, astrocytes, or co-cultures at higher transduction rates (50-90%). In particular, 40-60% of all motor neurons were successfully targeted by AAV5.2-GFP at a MOI = 1E06 (**Fig. 4b**). Transduction and expression of the transgene of interest (PC-OxPL-VecTab®) were correlated in a dose-dependent manner (**Fig. 4e**) as shown by the evaluation of PC-OxPL- VecTab® mRNA (**Fig. 4c**) and protein (**Fig. 4d**) content at a dose-dependent manner.

**Figure 4.**
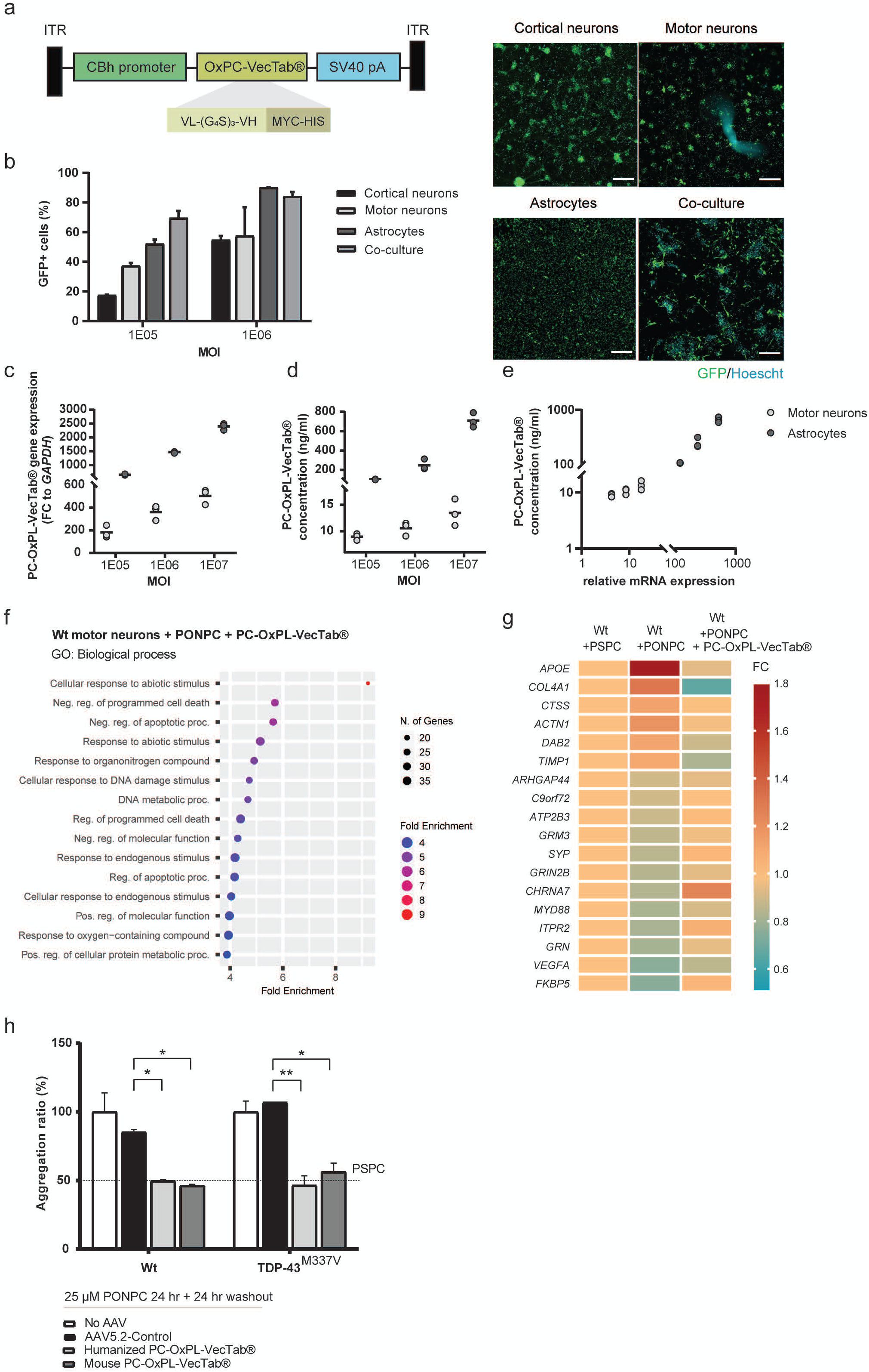
AAV-delivered PC-OxPL-VecTab®, targeting PC-OxPL, prevents neurotoxicity in iPSC-derived motor neurons. **a.** Schematic representation of AAV5.2-CBh- PC-OxPL-VecTab® cassette (ITR, inverted terminal repeats; CBh, CBA hybrid; SV40 pA, simian virus 40 PolyA). **b.** AAV5.2-GFP exhibits differential tropism among human-derived cells: cortical neurons, motor neurons, astrocytes, and motor neuronal:astrocytic co-culture. Values are expressed as means ± SEM; n = 3. Representative images are found on the *right panel* (GFP (FITC), green fluorescent protein). Scale bar = 300 µM. **c.** PC-OxPL-VecTab® transcript expression following AAV transduction of motor neurons and astrocytes. Both cell types were transduced with AAV5.2-Control at different MOIs, and cell lysate collected at eight days post-transduction. The scatter plot displays gene expression mean, normalized to *GAPDH* and relative to non-transduced condition (background); n = three. **d.** PC-OxPL- VecTab® protein expression following AAV transduction of motor neurons and astrocytes. Both cell types were transduced with AAV5.2-CBh-PC-OxPL-VecTab® at different MOIs and extracellular medium collected at eight days post-transduction. The scatter plot displays protein concentration. Maximum signal (OD450) using a 4x dilution was used to estimate scFv concentration; n = three. **e.** Correlation analysis of PC-OxPL-VecTab® transcript expression and protein concentration in motor neurons and astrocytes. Both cell types were transduced with AAV5.2-Control at different MOIs, and cell lysate collected at eight days post-transduction; n = three. **f.** Network analysis following AAV5.2- PC-OxPL-VecTab® transduction of wt motor neurons exposed to PC-OxPL (PONPC). Gene expression data from both Neuroinflammation and Neuropathology were used in the analysis. All pathways are sorted by Fold Enrichment value. **g.** Heatmap representation of transcriptome recovery following AAV5.2-Control transduction of wt motor neurons exposed to PC-OxPL (PONPC). **h.** TDP- 43 aggregation in wt and TDP-43^M337V^ motor neurons transduced with either mouse or humanized AAV5.2-PC-OxPL-VecTab® and exposed to PSPC or PC-OxPL. TDP-43 aggregation is shown as the ratio of aggregation measured by the HTRF TDP-43 assay. Values are expressed as means ± SEM, normalized to the assay positive control. Ordinary one-way ANOVA with Dunnett’s multiple comparison test, ***p < 0.001; n = 2 independent experiments, six replicates.

Next, we evaluated the efficacy of PC-OxPL-VecTab® in neutralizing the effects of PONPC toxicity in wt motor neurons. The transcripts influenced by PC-OxPL-VecTab® included molecular pathways such as regulation of cell death, apoptotic processes, protein metabolic processes, molecular function and response to endogenous stimulus (**Fig. 4f**). Approximately 50% of all PONPC-sensitive transcripts previously associated with ALS (**Fig. 2d**) (n = 18 out of 36) showed partial or complete restoration (≥0.1 FC) following treatment with PC-OxPL-VecTab® (**Fig. 4g**). Detailed information can be found in **Supplementary Table S7**.

We sought to determine the effect of PC-OxPL-VecTab® on ALS-associated phenotypes. To avoid artifacts or observations resulting from AAV transduction rather than specific AAV5.2-PC-OxPL- VecTab® effects, we first qualitatively evaluated the co-culture phenotypes under non-transduced and AAV5.2-Control conditions. Microscopic observation of the neuronal network of both wt and ALS motor neurons did not reveal any noticeable differences upon AAV5.2 transduction over time in culture (data not shown). This same AAV construct did not affect TDP-43 aggregation, as measured by the HTRF assay, in comparison to a non-transduced condition (**Fig. 4h**). Notably, we found that both mouse and humanized PC-OxPL-VecTab® were able to entirely prevent TDP-43 aggregation caused by PONPC exposure of wt or ALS motor neurons (**Fig. 4h**).

### PC-OxPL-VecTab® neutralizes PC-OxPL toxicity in a sALS CSF mouse model

While the evaluation of PC-OxPL-VecTab® *in vitro* offers a preliminary indication of its potential in neutralizing PC-OxPL toxicity, understanding its value in different models with added complexity is crucial from a therapeutic standpoint. To examine whether single dosing of AAV5.2-PC-OxPL- VecTab® could lead to *in vivo* functional improvements we selected the sALS CSF mouse model which has been shown to display several pathological features of ALS ^26^.

Adult female C57BL/6J wt mice were first injected with AAV5.2-PC-OxPL-VecTab® into the mid-cervical subarachnoid space. Four weeks after the AAV injection, the same mice were injected with sALS patient-derived CSF according to the same route of administration. We observed a moderate but clear increase in PC-OxPL signal in motor neurons following sALS CSF exposure (**Fig. 5a**). This was accompanied by the onset of functional disabilities, as shown by developed forelimb motor deficits and decreased grip strength, though the effects on the latter readout did not reach statistical significance (**Fig. 5b-c**), and motor neuron degeneration (**Fig. 5d**), as earlier described ^26^. Regarding neuroinflammatory phenotypes, a subtle increase was observed in GFAP and IBA1 expression in the spinal cords of sALS CSF-injected mice (**Supplementary Fig. S3a-b**).

**Figure 5.**
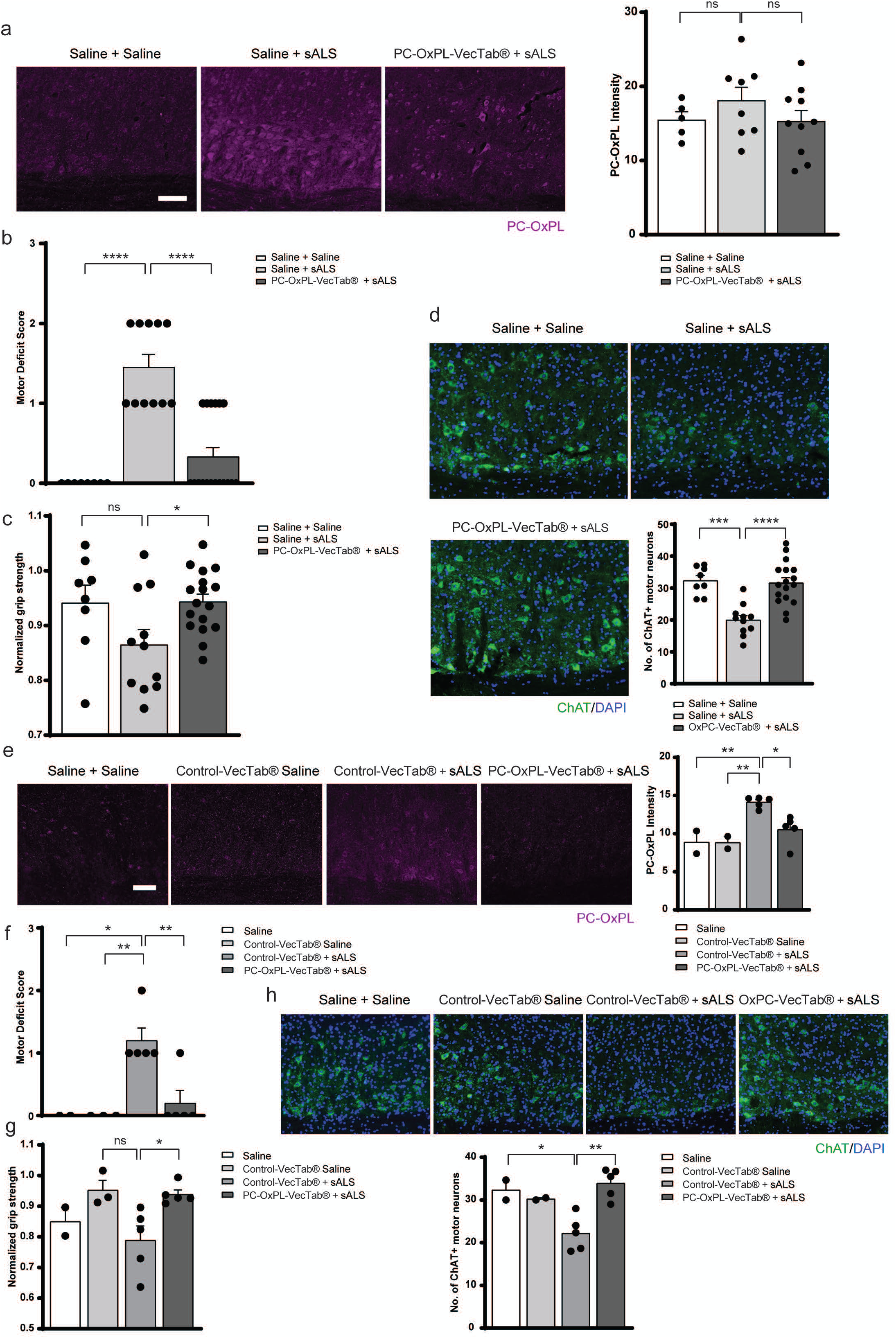
PC-OxPL-VecTab® attenuates sALS CSF-induced motor disability and motor neuron degeneration. **a.** *Left panel.* Representative images of PC-OxPL immunostaining in the cervical spinal cord at one day post intrathecal injection of saline or sALS CSF. Mice received intrathecal injections of saline or AAV5-2-PC-OxPL-VecTab® four weeks prior to sALS CSF injection. Scale bar = 100 µm. *Right panel.* Quantification of PC-OxPL immunostaining intensity in the ventral grey matter at one day post sALS/saline injection and four weeks days post AAV5.2-PC-OxPL-VecTab® injection. Values are expressed as means ± SEM. One-way ANOVA with Bonferroni’s test, ns, p ≥ 0.05. Saline + Saline (n = five mice), Saline + sALS (n = eight mice), AAV5.2-PC-OxPL-VecTab® + sALS (n = 10 mice). **b.** Motor deficit scores and **c.** Normalized forelimb grip strength force at one day post intrathecal injection of saline or sALS CSF. Mice received intrathecal injections of saline or AAV5.2-Control four weeks prior to sALS CSF injection. Values are expressed as means ± SEM. One-way ANOVA with Bonferroni’s test. One-way ANOVA with Bonferroni’s test, ****p < 0.0001, *p < 0.05. Saline + Saline (n = 11 mice), Saline + sALS (n = 11 mice), AAV5-2-PC-OxPL-VecTab® + sALS (n = 18 mice). **d.** *Upper panel.* Representative images of cervical spinal cords immunostained for ChAT at one day post intrathecal injection of saline or sALS CSF. *Lower panel.* Quantification of the number of ChAT+ motor neurons in cervical ventral horns at one day post sALS/saline injection and four weeks post AAV5.2-PC-OxPL-VecTab® injection. Mice received intrathecal injections of saline or AAV5.2- PC- OxPL-VecTab® four weeks prior to sALS CSF injection. One-way ANOVA with Bonferroni’s test, ****p < 0.0001, ***p < 0.001. Saline + Saline (n = 11 mice), Saline + sALS (n = 11 mice), AAV5-2- PC-OxPL-VecTab + sALS (n = 18 mice). **g.** Representative images of PC-OxPL immunostaining in the cervical spinal cord at one day post intrathecal injection of Saline or sALS CSF, previously incubated with PC-OxPL-VecTab®. Scale bar = 100 µm. **e.** Quantification of PC-OxPL immunostaining intensity in the ventral grey matter after intrathecal injection of Saline or sALS CSF, previously incubated with PC-OxPL-VecTab® or Control-VecTab®. Values are expressed as means ± SEM. One-way ANOVA with Bonferroni’s test, **p < 0.01, *p < 0.05. Saline/Control-VecTab® + Saline (n = two mice), Control-VecTab® + sALS/PC-OxPL-VecTab® + sALS (n = five mice). **f.** Motor deficit scores and **g.** Normalized forelimb grip strength force after intrathecal injection of saline or sALS CSF, previously incubated with PC-OxPL-VecTab® or Control-VecTab®. Values are expressed as means ± SEM. One- way ANOVA with Bonferroni’s test, **p < 0.01, *p < 0.05. Saline (n = two), Control-VecTab® + Saline (n = three mice), Control-VecTab® + sALS/ PC-OxPL-VecTab® + sALS (n = five mice). **h.** *Upper panel.* Representative images of cervical spinal cords immunostained for ChAT after intrathecal injection of saline or sALS CSF, previously incubated with PC-OxPL-VecTab® or Control-VecTab®. Scale bar = 100 µm. *Lower panel.* Quantification of the number of ChAT+ motor neurons in cervical ventral horns after intrathecal injection of saline or sALS CSF, previously incubated with PC-OxPL- VecTab® or Control-VecTab®. Values are expressed as means ± SEM. One-way ANOVA with Bonferroni’s test, **p < 0.01, *p < 0.05. Saline (n = two mice), Control-VecTab® + Saline (n = two mice), Control-VecTab® + sALS/PC-OxPL-VecTab® + sALS (n = five mice).

In parallel, we examined the protective effect of AAV5.2-PC-OxPL-VecTab® administration in reverting these phenotypes. Intrathecal injection with AAV5.2-PC-OxPL-VecTab® and subsequent sALS CSF led to a normalization of PC-OxPL levels in the spinal cord (**Fig. 5a**). At the same time, it prevented motor impairments, as shown by the motor deficit scores and grip strength levels comparable to those injected with saline (**Fig. 5b-c**). Furthermore, it completely abolished motor neuron death caused by sALS CSF injection, as shown by a similar number of ChAT+ motor neurons as saline-injected mice (**Fig. 5d**). We assessed whether AAV5.2-PC-OxPL-VecTab® would also be efficient in protecting against the minor neuro-inflammatory changes present in the sALS CSF-injected mice. Despite the mild differences between saline and sALS CSF-injected mice, AAV5.2-PC-OxPL-VecTab normalized GFAP and IBA1 expression towards saline levels (**Supplementary Fig. S3a-b**).

Thus far, the results described underscore a link between PC-OxPL and CSF-mediated toxicity in ALS. However, these findings do not rule out the possibility that sALS CSF toxicity may result from phospholipid oxidation products (e.g., 4’-HNE or MDA)^14^ rather than direct PC-OxPL toxicity. Hence, we investigated the effect of purified non-vectorized PC-OxPL-VecTab® preincubation with sALS CSF prior to intrathecal injection. This approach revealed that preincubation of sALS CSF with PC- OxPL-VecTab®, but not with Control-VecTab®, prevented overall sALS CSF toxicity, including PC- OxPL accumulation (**Fig. 5e**), motor deficits (**Fig. 5f**), grip strength decline (**Fig. 5g**), and motor neuron degeneration (**Fig. 5h**). Also, preincubation with PC-OxPL-VecTab® prevented TDP-43 mislocalization induced by sALS CSF treatment, however, the effects were not statistically significant (**Supplementary Fig. S3c**). No differences were found in GFAP/IBA1 expression upon sALS CSF preincubation with either Control- or PC-OxPL-VecTab® (**Supplementary Fig. S3d-e**).

Altogether, these findings demonstrate that PC-OxPL are key contributors to sALS pathology via CSF transfer and that PC-OxPL-VecTab® effectively reverses this toxicity.

### Widespread biodistribution in brain and spinal cord and safe expression profile of PC-OxPL- VecTab® in mini pigs

Having established the efficacy of PC-OxPL-VecTab® in neutralizing PC-OxPL toxicity using *in vitro* and *in vivo* approaches, we sought to determine its therapeutic feasibility.

Due to large differences in anatomy, brain size, and spinal cord length between rodents and humans, studies in large animals are key to establishing a clinical development trajectory. In ALS, the spinal cord and motor cortex are the primary affected regions and, upon intrathecal injection, the CSF flow contributes to the distribution of the AAV5.2 vector along the spinal cord and cortical regions (**Fig. 6a**).

**Figure 6.**
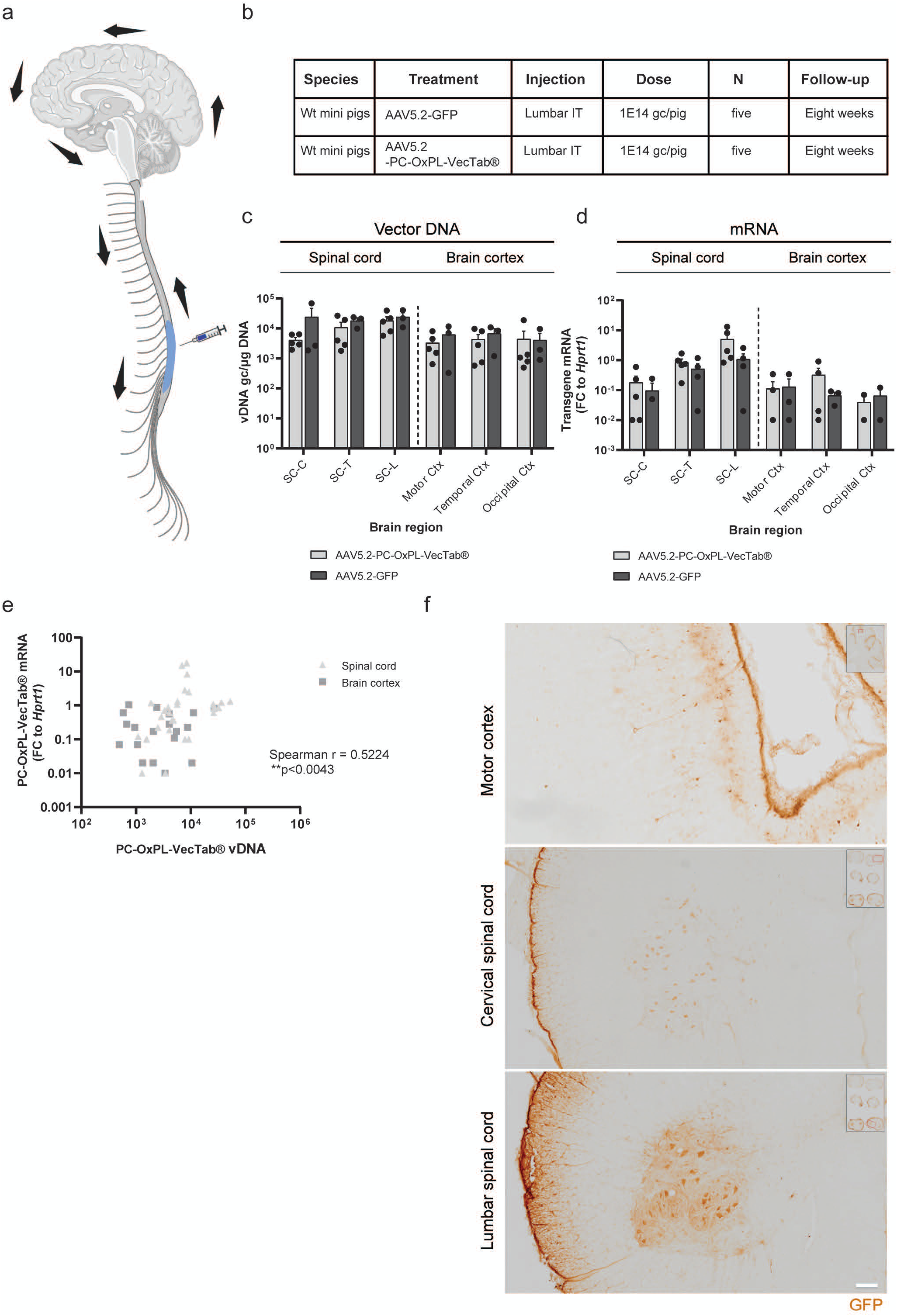
Widespread biodistribution of PC-OxPL-VecTab® in brain and spinal cord in mini pigs upon intrathecal administration. **a.** Schematic representation of CSF flow (arrows) and biodistribution of AAV in the different CNS areas upon intrathecal administration. **b.** Study design and treatment groups of intrathecal delivery of AAV-GFP and AAV5.2-Control in wt mini pigs. **c.** Vector DNA levels (genome copies/ug DNA) in the spinal cord and brain cortex at eight weeks after AAV- intrathecal delivery. **d.** Transgene mRNA levels (FC to housekeeping gene *Hprt1*) of GFP or PC-OxPL- VecTab® in the spinal cord and brain cortex of mini pigs eight weeks after AAV-intrathecal delivery. **e.** Correlation analysis of vector DNA and PC-OxPL-VecTab® mRNA expression in mini pig’s spinal cord and brain cortex. Spearman correlation analysis, r = 0.5224, **p = 0.0043. **f.** Detection of GFP protein by immunohistochemistry in the motor cortex *(top),* cervical spinal cord *(center)* and lumbar spinal cord *(bottom)* of mini pigs IT-treated with AAV-GFP.

To investigate the feasibility of intrathecal injection to target the spinal cord and motor cortex, we investigated the biodistribution and expression level of AAV5.2-PC-OxPL-VecTab® in the CNS of large animals upon a one-time administration. Five wt mini pigs were injected with AAV5.2-PC-OxPL- VecTab® or AAV5.2-GFP via intrathecal administration in the lumbar region of the spinal cord (**Fig. 6b**). After injection, the animals were followed for eight weeks before necropsy. The biodistribution of AAV5.2 was determined by vector DNA (vDNA) genome copies (gc) in the different segments of the spinal cord (cervical, thoracic and lumbar), in the motor cortex, temporal cortex and occipital cortex. Consistent with the route of injection and CSF flow, high levels of vDNA were detected in the spinal cord, being the highest in the lumbar region, and in all cortical areas (**Fig. 6c**). AAV5.2-PC-OxPL- VecTab® and -GFP mRNA expression levels were measured in the same regions and quantified as FC to the porcine housekeeping gene *Hprt1* (**Fig. 6d**). The highest levels of transgene expression (>1 FC to *Hprt1)* were detected in the lumbar spinal cord. Correlation analysis between vector DNA levels and mRNA levels showed a significant positive correlation between transduction levels and transgene expression in both spinal cord and brain cortex tissues (**Fig. 6e**). To visualize the AAV distribution in the brain, GFP immunohistochemical analysis was performed. The motor cortex, particularly large pyramidal neurons in layer V, showed GFP signal (**Fig. 6f**). In the cervical and in lumbar spinal cord, a high number of large cells in the ventral horn, potentially motor neurons, were positive for GFP staining. Taken together, these results show that the intrathecal administration of AAV5.2-PC-OxPL-VecTab® is an effective route of administration to target the primary ALS-affected areas - spinal cord and cortex -, in large animals.

## Discussion

This study provides several levels of evidence that PC-OxPL are associated with the pathology present in motor neurons in the brain and spinal cord in patients with ALS. PC-OxPL levels on APOE were enriched in the in the CSF relative to plasma suggesting local generation. Exposure of PC-OxPL to iPSC-derived motor neurons recapitulated the transcriptome changes and TDP-43 pathology known to be associated with ALS. These effects were ameliorated *in vitro* and *in vivo* by overexpression of a well-described antibody fragment (PC-OxPL-VecTab®), suggesting that sequestering PC-OxPL may prevent ALS pathology. These findings provide a rationale to develop PC-OxPL targeted therapies to improve the prognosis of patients with sALS.

TDP-43 proteinopathy is a primary hallmark of sALS, driving extensive mis-splicing, the formation of “cryptic” transcripts and peptides, and the loss of key proteins essential for motor neuron homeostasis. Further, TDP-43 aggregation impairs axonal transport and disrupts neuromuscular synapses eventually causing detrimental effects on motor neurons ^4,42^. While oxidative stress and aging have been proposed to contribute to motor neuron degeneration in ALS, the mechanisms underlying TDP-43 misfolding and aggregation remain elusive ^4,50,51^. Ferroptosis, a programmed cell death pathway characterized by PUFA oxidation, has gained recognition as a mechanism contributing to selective motor death in ALS ^8,9^. PC- OxPL accumulation is tightly linked to ferroptosis and their association with multiple pathologies has sparked interest in the potential role of PC-OxPL in neurodegenerative diseases, such as ALS ^13,18,32^.

Our study identified PC-OxPL accumulation in the most affected CNS regions in ALS, including those mirroring the spread of TDP-43 pathology ^27^. This is the first evidence linking PC-OxPL to TDP-43 proteinopathy and ALS pathology. PC-OxPL accumulation was predominantly associated with motor neurons in ALS spinal cord samples and reflected by elevated PC-OxPL levels in CSF from ALS patients compared to HC. Interestingly, the PC-OxPL signatures in ALS CSF differed from those observed in multiple sclerosis (MS) ^19^, possibly reflecting differences in disease-specific processes ^18,52^. Also, PC abundance is somewhat tissue-specific, hence, it is likely that PC-OxPL generation is not a stochastic event ^18,19,29^. Given the distinct profiles in different pathologies, these findings suggest that specific PC-OxPL may contribute to the unique pathophysiology of various neurodegenerative conditions.

PC-OxPL in CSF, but not in plasma, was primarily associated with APOE in healthy and ALS-affected brains. Unlike plasma, where a variety of lipoproteins are present, CSF mainly contains APOE particles (reviewed in ^53^). These particles are locally produced and reflect lipid changes within the CNS, due to the restrictive nature of the blood-brain barrier. Notably, only APOA-I particles are able to cross this barrier ^54^. The differential distribution of PC-OxPL in apolipoproteins in brain in comparison to the periphery is suggestive of specific OxPL metabolism in the CNS, including locally restricted generation and distribution within CNS regions and cell types. With the current methods, it is not possible to ascertain if a specific APOE isoform was most associated with OxPL binding, if the APOE particles contain lipids or where the OxPL may be present - if non-covalently bound in the lipid phase or covalently bound to APOE amino acids. In prior studies, it was demonstrated that OxPL can be present both in the LDL-like moiety of Lp(a) as well as covalently bound to the apolipoprotein(a) component ^55,56^. Glial cells are the main source of CNS APOE ^57,58^, but under pathological conditions, neurons can also secrete APOE, increasing the ability for lipid transfer to astrocytes ^44,59^. Notably, *APOE* was the most upregulated gene in motor neurons exposed to PONPC. It has been recently reported that APOE isoforms differ in their capacity to transfer cholesteryl-PUFA esters to neuronal cells, related to their affinity for the LDL receptor, APOE4 having the highest affinity and APOE2 the lowest ^60^. Conversely, TDP-43 pathology is more severe in mice with a human *APOE2* background as compared to other isoforms ^61^. The relationship between the onset or progression of ALS and APOE isoforms is inconclusive. Still, it is tempting to speculate that the acute upregulation of *APOE* following exposure to PC-OxPL constitutes a protective response of diseased neurons to excessive fatty acid (FA) oxidation. The mechanism by which primary astrocytes metabolize excessive neuronal FA contained in APOE particles via β-mitochondrial oxidation has been described ^44^. However, it is also possible that APOE particles carrying PC-OxPL induce or exacerbate motor neuron toxicity when taken up by receptors such as low-density lipoprotein receptors (LDLR or LRP) ^62^.

Previous work has shown that intrathecal transfer of sALS CSF into wt mice induces profound motor neuron loss, TDP-43 mislocalization, and motor dysfunction ^26^. These effects has been observed using sALS CSF but not following the injection of CSF from familial ALS (fALS) patients with *SOD1*, *C9orf72*, or *TARDBP* mutations ^26^. In this study, we used a previously characterized sALS CSF sample that elicited motor disability and histochemical signatures associated with ALS ^26^. Remarkably, the observed toxicity markedly affected the degeneration of ChAT+ motor neurons and was accompanied by TDP-43 mislocalization, suggesting that these neurons are particularly susceptible to PC-OxPL- induced damage via sALS CSF transfer. TDP-43 regulates thousands of transcripts, most of which encode proteins involved in synaptic homeostasis (e.g. *STMN2*, *UNC13A*), and this has been shown to underline TDP-43 loss-of-function ^38^. Notably, intrathecal injection of mice with PC-OxPL-VecTab® reduced PC-OxPL levels in the CNS, prevented TDP-43 mislocalization, and protected motor neurons from degeneration. As a result, grip strength and motor deficit scores were substantially improved in treated mice. To confirm that PC-OxPL were the driver of the observed pathology, CSF samples were pretreated with an PC-OxPL-binding antibody before transfer. Remarkably, this treatment effectively prevented TDP-43 mislocalization, motor neuron degeneration and the onset of motor deficits. These findings suggest that the presence of PC-OxPL in the CSF of sALS patients drives significant TDP-43 pathology and motor neuron degeneration. We show that these effects can be mitigated by a therapeutic scFv that targets the PC-OxPL neoepitope formed through lipid oxidation. These observations fit well with prior evidence demonstrating that POVPC injection induced spinal cord demyelination in mice hence that PC-OxPL species mediate neurodegeneration in vivo ^19^. Intriguingly, previous work by Wong *et al.* has shown that APOB mediates the spread of sALS pathology using sALS CSF mouse model ^26^. At first, this may appear contradictory to our findings suggesting APOE as the main contributor to PC-OxPL neurotoxicity in ALS. It could indicate a more negligible contribution of APOB to PC-OxPL transport and neurotoxicity which is in line with PC-OxPL detection on APOB particles in ALS CSF. Additionally, challenges may arise in replicating APOE-PC-OxPL interactions under non-physiological conditions or in cases of specific APOB depletion, which could account for the discrepancies between the current and the previous study ^26^.

Our findings warrant further investigation into several critical aspects. The exact mechanism by which PC-OxPL mediates TDP-43 proteinopathy and eventual motor neuron degeneration remains elusive. Others have demonstrated that TDP-43 (in)solubility and misfolding are regulated by cysteine oxidation and disulfide bond formation (reviewed in ^4^). More recently, it was reported that TDP-43 contains a domain similar to the FERM, ARH/RhoGEF And Pleckstrin Domain Protein 1 (FARP1) lipid-binding domain and that exposure of TDP-43 to cholesterol or PC induced *in vitro* fibrillation of recombinant TDP-43 constructs ^63^. Hence, the binding of TDP-43 to PC may directly be involved in TDP-43 aggregation, and it would be interesting to compare the effects of the PC-OxPL species found in the current study on TDP-43 aggregation in more detail. On a general note, the fact that PC-OxPL may be a contributing mechanism to another underlying process in ALS cannot be ruled out. As far as the field is concerned, it is plausible to assume that PC-OxPL may act as mediators of general oxidative stress or, in particular, ferroptosis-associated mechanisms ^9,18,64^. Further studies are also required to assess if the PC-OxPL are simply inactivated or are additionally cleared as a mechanism of benefit, as suggested in other CNS pathologies ^19^. Nonetheless, the benefits of targeting PC-OxPL neurotoxicity in ALS are evident and encouraging.

The success of a gene therapy-based construct is dependent mainly on its ability to reach regions of interest. Building on the remarkable efficacy of PC-OxPL-VecTab® in the sALS CSF mouse model, we explored its potential therapeutic application in humans by evaluating its biodistribution and transgene expression in mini pigs. After intrathecal administration, widespread transgene expression was observed in the spinal cord and motor cortex. A GFP-expressing construct was delivered with the same capsid, demonstrating tropism in regions of interest. The treatment was well-tolerated, and no adverse effects were observed during an eight-week follow-up.

In summary, our work provides the first evidence that PC-OxPL accumulation is an important element in ALS pathology. We describe a novel therapeutic approach using an AAV-delivered PC-OxPL- VecTab®, which effectively neutralizes PC-OxPL neurotoxicity both *in vitro* and in *vivo*. Finally, we establish clinical relevance by demonstrating a favorable distribution and safety profile in a large animal. This study positions PC-OxPL as an important neurotoxic factor in ALS and highlights PC- OxPL-VecTab® as a promising candidate for innovative ALS therapies. Moreover, our findings suggest that PC-OxPL-VecTab® could be extended to other lipid-related diseases where PC-OxPL toxicity plays a significant role.

## Abbreviations and symbols

4-HNE: 4-hydroxynonenal
AAV: Adeno-associated virus
ALS: Amyotrophic Lateral Sclerosis
ApoB: Apolipoprotein B
ApoE/APOE: Apolipoprotein E
AD: Alzheimer’s Disease
BSA: Bovine serum albumin
CBh: Chicken β-actin hybrid
CSF: Cerebrospinal fluid
ChAT: Choline Acetyltransferase
DAPI: 4′,6-diamidino-2-phenylindole
ETOH: Ethanol
FA: Fatty acids
fALS: familial Amyotrophic Lateral Sclerosis
FARP1: FERM, ARH/RhoGEF And Pleckstrin Domain Protein 1
FC: Fold-change
FDR: False discovery rate
FOV: Fields of view
GO: Gene ontology
GFAP: Glial fibrillary acidic protein
H: hour(s)
hE06: Humanized E06 protein
IBA1: Ionized Calcium Binding Adaptor Molecule 1
iPSC: Induced pluripotent stem cells
LDL: Low-density lipoproteins
M: minute(s)
MDA: Malondialdehyde
mE06: Mouse E06
MNP: Motor neuron progenitors
MS: Multiple sclerosis
MOI: Multiplicity of infection
mRNA: Messenger RNA
NBB: Netherlands Brain Bank
OxLDL: Oxidized LDL
OxPL: Oxidized phospholipids
PBS: Phosphate-buffered saline
PC: Phosphatidylcholine
PC-OxPL: PC-containing oxidized phospholipids
PD: Parkinson’s Disease
PDL: Poly-D-Lysine
PONPC: *1-palmitoyl-2-(9-oxononanayl)-phosphocholine*
PSPC: *1-hexadecanoyl-2-octadecanoyl-sn-glycero-3-phosphocholine*
PUFA: Poly-unsaturated fatty acids
pTDP-43: Phosphorylated TDP-43
QC: Quality control
ROS: Reactive oxygen species
Rpm: rotations per minute
rRNA: Ribosomal RNA
RT: Room temperature
S: seconds
sALS: Sporadic Amyotrophic Lateral Sclerosis
scFv: Single-chain variable fragment
SD: Standard deviation
SEM: Standard error of the mean
SOD1: Superoxide dismutase 1
TARDBP: Transacting response DNA-binding protein
TBS: Tris-buffered saline
TDP-43: TAR DNA-binding protein
TUBB3: βIII Tubulin

## Data availability

The datasets used (NanoString nCounter profiling) are deposited in the Gene Expression Omnibus (GEO) database under the accession numbers GSE281698 and GSE281714.

## Acknowledgments

The sALS animal model studies performed at Tisch MSRCNY were funded by the Laurence and Sandi Gluck Charitable Foundation.

## Author contributions

SVD and AG-D led the project. Experiments were designed and performed by AG-D, JKW, SP-V, WP, RH, MS-G, SP, RVD, FS, MG and ST, and data interpretation and analysis conducted by AG-D, JKW, SP-V, WP, RH, MS-G, SP, MG and ST. Experimental work performed by AG-D, SP-V, WP, RH, MS- G, SP, RVD and FS was supervised by PSK and SVD. Experimental work conducted by JKW was supervised by SAS. SVD and AG-D wrote the first draft of the manuscript, and all authors contributed by writing/editing the manuscript until its final form.

## Supplementary data

**Supplementary Figure S1. a.** Morphological representation of Axol motor neurons exposed to frozen- or freshly prepared lipid species (PSPC, PONPC and PaZPC) at a dose range (25, 50 and 100 µM) at 24 h post PC-OxPL exposure, based on Calcein TM (Invitrogen) live imaging. Displayed images were selected from a panel of six replicates per condition. Scale bar = 100 µm. **b.** % of PC-OxPL+ cells following wt motor neurons exposure to PONPC (25 µM). Values are expressed as means ± SEM. Two-way ANOVA, ns, p ≥ 0.05; n= two independent experiments, six replicates. Scale bar = 100 µm.

**Supplementary Figure S2. a.** TDP-43 aggregation in wt and TDP-43^M337V^ motor neurons exposed to 0.1% DMSO or 0.1 µM of MG-132. TDP-43 aggregation is shown as the ratio of aggregation measured by the HTRF TDP-43 assay. Values are expressed as means ± SEM, normalized to the assay positive control. Ordinary one-way ANOVA with Tukey’s multiple comparison test, ***p < 0.001; n = two independent experiments, six replicates. **b.** pTDP-43 expression following PSPC or PONPC exposure of wt Axol motor neurons. Values are expressed as means ± SEM, normalized to the control condition (24 h 25 µM PSPC + 6 or 24 h washout). Two-way ANOVA, ns, p ≥ 0.05; n= two independent experiments, 12 replicates. Scale bar = 100 µm. Alterations on mitochondrial profile following wt motor neurons exposure to PONPC, including **c.** % of mitochondria+ cells and **d.** mitochondria expression, as assessed with immunostaining using mouse anti-mitochondrial (ab92824) antibody. Values are expressed as means ± SEM, normalized to the control condition (24 h 25 µM PSPC). **e.** PC-OxPL levels as detected on the surface of different apolipoprotein particles as measured in paired CSF from HC and ALS patients. Values are expressed as means ± SEM; each dot represents a single individual (n = 15).

**Supplementary Figure S3. a.** *Left panel.* Representative images of GFAP immunostaining in the cervical spinal cord at one day post intrathecal injection of saline or sALS CSF. Mice received intrathecal injections of saline or AAV5.2-PC-OxPL-VecTab® four weeks prior to sALS CSF injection. Scale bar = 100 µm. *Right panel.* Quantification of GFAP immunostaining intensity in the dorsal white matter at one day post sALS/saline injection and four weeks post AAV5.2-PC-OxPL-VecTab® injection. Values are expressed as means ± SEM. One-way ANOVA with Bonferroni’s test, ns, p ≥ 0.05. **b.** *Left panel.* Representative images of IBA1 immunostaining in the cervical spinal cord at one day post intrathecal injection of saline or sALS CSF. Mice received intrathecal injections of saline or AAV5.2-PC-OxPL-VecTab® four weeks prior to sALS CSF injection. Scale bar = 100 µm. *Right panel.* Quantification of IBA1 immunostaining intensity in the ventral grey matter at one day post sALS/saline injection and four weeks post AAV5.2-PC-OxPL-VecTab® injection. Values are expressed as means ± SEM. One-way ANOVA with Bonferroni’s test, ns, p ≥ 0.05. **c.** *Left panel.* Representative images of TDP-43 signal on ChAT+ neurons in cervical spinal cords following intrathecal injection of saline or sALS CSF, previously incubated with PC-OxPL-VecTab® or Control-VecTab®. *Right panel.* Quantification of the % of ChAT+ neurons expressing cytoplasmic TDP-43 following intrathecal injection of saline or sALS CSF, previously incubated with PC-OxPL-VecTab® or Control-VecTab®. Values are expressed as means ± SEM. One-way ANOVA with Bonferroni’s test, ns, p ≥ 0.05. **d.** *Upper panel.* Representative images of GFAP immunostaining in cervical spinal cords following intrathecal injection of saline or sALS CSF, previously incubated with PC-OxPL-VecTab® or Control-VecTab®. *Lower panel.* Quantification of GFAP immunostaining following intrathecal injection of Saline or sALS CSF, previously incubated with PC-OxPL-VecTab® or Control-VecTab®. Values are expressed as means ± SEM**. e.** *Upper panel.* Representative images of IBA1 immunostaining in cervical spinal cords following intrathecal injection of saline or sALS CSF, previously incubated with PC-OxPL- VecTab® or Control-VecTab®. *Lower panel.* Quantification of IBA1 immunostaining following intrathecal injection of saline or sALS CSF, previously incubated with PC-OxPL-VecTab® or Control- VecTab®. Values are expressed as means ± SEM.

**Supplementary Table S1. Details on the sample selection and demographics of the human samples used in this study** (NDC, non-demented control, f, female; m, male; PMD, postmortem delay; NBB, Netherlands Brain Bank; NA, non-applicable; SD, standard deviation; HC, healthy control).

Supplementary Table S2. List of primary and secondary antibodies used in the various studies (ON, overnight; ICC, immunocytochemistry; IHC, immunohistochemistry).

**Supplementary Table S3.** List of monitored PC-OxPL by LC-MS for targeted analysis in CSF and plasma with the indication in which matrix they were detected.

**Supplementary Table S4. List of differently expressed (DE) transcripts in ALS (TDP-43^M337V^ and SOD1^G93A^) and wt + PONPC motor neurons, including overlapping changes, as assessed by NanoString nCounter analysis.** <u>Bioinformatics and Systems Biology (ugent.be)</u> was used in the construction of Venn Diagrams (DE, differently expressed; NP, neuropathology, NI neuroinflammation; &, overlapping) Supplementary Table S5. List of overlapping differently expressed (DE) transcripts between wt + PONPC motor neurons and ALS/TDP-43 knockdown (KD) databases**. Data mining analysis was performed using generated NanoString data and three independent available datasets: (1) ALSoD** ^33^**, (2) Postmortem spinal cord tissue from patients with sALS** ^34^ **and (3) iPSC-derived neurons with reduced TDP-43 expression following CRISPRi** ^37^ **(&, overlapping).**

**Supplementary Table S6. List of differently expressed (DE) transcripts with an association with ferroptosis.** Association of (1) DE transcripts following PONPC exposure of wt motor neurons and (2) with an overlap to ALS with ferroptosis. FerrDb was used as a database dedicated to gene associations with ferroptosis ^40^ (&, overlapping).

**Supplementary Table S7. List of PONPC-sensitive transcripts following transduction with AAV5.2-PC-OxPL-VecTab®.**Data is expressed as FC normalized to the physiological conditions (Wt+ PSPC).

## References

1. Chiò, A. et al. Prognostic factors in ALS: A critical review. Amyotrophic Lateral Sclerosis 10, 310–323 (2009).

2. Hardiman, O. et al. Amyotrophic lateral sclerosis. Nat Rev Dis Primers 3, (2017).

3. 3. Jaiswal, M. K. Riluzole and edaravone: A tale of two amyotrophic lateral sclerosis drugs. Medicinal Research Reviews vol. 39 733–748 Preprint at 10.1002/med.21528 (2019).

4. Prasad, A., Bharathi, V., Sivalingam, V., Girdhar, A. & Patel, B. K. Molecular mechanisms of TDP-43 misfolding and pathology in amyotrophic lateral sclerosis. Front Mol Neurosci 12, (2019).

5. Donde, A. et al. Splicing repression is a major function of TDP-43 in motor neurons. Acta Neuropathol 138, 813–826 (2019).

6. Moujalled, D., Strasser, A. & Liddell, J. R. Molecular mechanisms of cell death in neurological diseases. Cell Death Differ 28, 2029–2044 (2021).

7. Wang, L. Y. et al. The Role of Ferroptosis in Amyotrophic Lateral Sclerosis Treatment. Neurochemical Research vol. 49 2653–2667 Preprint at 10.1007/s11064-024-04194-w (2024).

8. Wang, T. et al. Ferroptosis mediates selective motor neuron death in amyotrophic lateral sclerosis. Cell Death Differ 29, 1187–1198 (2022).

9. Yan, H. fa et al. Ferroptosis: mechanisms and links with diseases. Signal Transduct Target Ther 6, (2021).

10. Lee, H. et al. Multi-omic analysis of selectively vulnerable motor neuron subtypes implicates altered lipid metabolism in ALS. Nat Neurosci 24, 1673–1685 (2021).

11. Blasco, H. et al. Lipidomics Reveals Cerebrospinal-Fluid Signatures of ALS. Sci Rep 7, (2017).

12. Beers, D. R. et al. Tregs Attenuate Peripheral Oxidative Stress and Acute Phase Proteins in ALS. Ann Neurol 92, 195–200 (2022).

13. Muñoz, U. et al. Main Role of Antibodies in Demyelination and Axonal Damage in Multiple Sclerosis. Cell Mol Neurobiol 42, 1809–1827 (2022).

14. Binder, C. J., Papac-Milicevic, N. & Witztum, J. L. Innate sensing of oxidation-specific epitopes in health and disease. Nat Rev Immunol 16, 485–497 (2016).

15. Sun, X. et al. Neutralization of Oxidized Phospholipids Ameliorates Non-alcoholic Steatohepatitis. Cell Metab 31, 189–206.e8 (2020).

16. Que, X. et al. Oxidized phospholipids are proinflammatory and proatherogenic in hypercholesterolaemic mice. Nature 558, 301–306 (2018).

17. Tsimikas, S. & Witztum, J. L. Oxidized phospholipids in cardiovascular disease. Nature Reviews Cardiology vol. 21 170–191 Preprint at 10.1038/s41569-023-00937-4 (2024).

18. Dong, Y. & Yong, V. W. Oxidized phospholipids as novel mediators of neurodegeneration. Trends Neurosci 45, 419–429 (2022).

19. Dong, Y. et al. Oxidized phosphatidylcholines found in multiple sclerosis lesions mediate neurodegeneration and are neutralized by microglia. Nat Neurosci 24, 489–503 (2021).

20. Tsimikas, S. et al. Lipoprotein(a) Reduction in Persons with Cardiovascular Disease. New England Journal of Medicine 382, 244–255 (2020).

21. Arai, K. et al. Acute impact of apheresis on oxidized phospholipids in patients with familial hypercholesterolemia. J Lipid Res 53, 1670–1678 (2012).

22. Giera, M. Clinical Metabolomics Methods and Protocols Second Edition Methods in Molecular Biology 2855.

23. Que, X. et al. Oxidized phospholipids are proinflammatory and proatherogenic in hypercholesterolaemic mice. Nature 558, 301–306 (2018).

24. Chen, H. Intron splicing-mediated expression of AAV rep and cap genes and production of AAV vectors in insect cells. Molecular Therapy 16, 924–930 (2008).

25. Xijin Ge, S., Jung, D. & Yao, R. ShinyGO: a graphical gene-set enrichment tool for animals and plants. doi:10.5281/zenodo.1451847.

26. Wong, J. K. et al. Apolipoprotein B-100-mediated motor neuron degeneration in sporadic amyotrophic lateral sclerosis. Brain Commun 4, (2022).

27. Kawakami, I., Arai, T. & Hasegawa, M. The basis of clinicopathological heterogeneity in TDP- 43 proteinopathy. Acta Neuropathologica vol. 138 751–770 Preprint at 10.1007/s00401-019-02077-x (2019).

28. Cortie, C. H. et al. Of mice, pigs and humans: An analysis of mitochondrial phospholipids from mammals with very different maximal lifespans. Exp Gerontol 70, 135–143 (2015).

29. Choi, J. et al. Comprehensive analysis of phospholipids in the brain, heart, kidney, and liver: brain phospholipids are least enriched with polyunsaturated fatty acids. Mol Cell Biochem 442, 187–201 (2018).

30. Hancock, S. E., Friedrich, M. G., Mitchell, T. W., Truscott, R. J. W. & Else, P. L. The phospholipid composition of the human entorhinal cortex remains relatively stable over 80 years of adult aging. Geroscience 39, 73–82 (2017).

31. Hawrot, J., Imhof, S. & Wainger, B. J. Modeling cell-autonomous motor neuron phenotypes in ALS using iPSCs. Neurobiol Dis 134, (2020).

32. Greig, F. H., Kennedy, S. & Spickett, C. M. Physiological effects of oxidized phospholipids and their cellular signaling mechanisms in inflammation. Free Radic Biol Med 52, 266–280 (2012).

33. Abel, O., Powell, J. F., Andersen, P. M. & Al-Chalabi, A. ALSoD: A user-friendly online bioinformatics tool for amyotrophic lateral sclerosis genetics. Hum Mutat 33, 1345–1351 (2012).

34. D’Erchia, A. M. et al. Massive transcriptome sequencing of human spinal cord tissues provides new insights into motor neuron degeneration in ALS. Sci Rep 7, (2017).

35. Uhlen, M. et al. A genome-wide transcriptomic analysis of protein-coding genes in human blood cells. Science (1979) 366, (2019).

36. Hayes, L. R. & Kalab, P. Emerging Therapies and Novel Targets for TDP-43 Proteinopathy in ALS/FTD. Neurotherapeutics vol. 19 1061–1084 Preprint at 10.1007/s13311-022-01260-5 (2022).

37. Brown, A. L. et al. TDP-43 loss and ALS-risk SNPs drive mis-splicing and depletion of UNC13A. Nature 603, 131–137 (2022).

38. Willemse, S. W. et al. UNC13A in amyotrophic lateral sclerosis: From genetic association to therapeutic target. *Journal of Neurology*, Neurosurgery and Psychiatry vol. 94 649–656 Preprint at 10.1136/jnnp-2022-330504 (2023).

39. Mohan, S. et al. Role of ferroptosis pathways in neuroinflammation and neurological disorders: From pathogenesis to treatment. Heliyon 10, (2024).

40. Zhou, N. & Bao, J. FerrDb: A manually curated resource for regulators and markers of ferroptosis and ferroptosis-disease associations. Database 2020, (2020).

41. Watanabe, S. et al. ALS-linked TDP-43M337V knock-in mice exhibit splicing deregulation without neurodegeneration. Mol Brain 13, (2020).

42. Scotter, E. L., Chen, H. J. & Shaw, C. E. TDP-43 Proteinopathy and ALS: Insights into Disease Mechanisms and Therapeutic Targets. Neurotherapeutics 12, 352–363 (2015).

43. Schönfeld, P. & Reiser, G. Why does brain metabolism not favor burning of fatty acids to provide energy-Reflections on disadvantages of the use of free fatty acids as fuel for brain. Journal of Cerebral Blood Flow and Metabolism 33, 1493–1499 (2013).

44. Ioannou, M. S. et al. Neuron-Astrocyte Metabolic Coupling Protects against Activity-Induced Fatty Acid Toxicity. Cell 177, 1522–1535.e14 (2019).

45. Mahley, R. W. Central nervous system lipoproteins: ApoE and regulation of cholesterol metabolism. Arterioscler Thromb Vasc Biol 36, 1305–1315 (2016).

46. Itabe, H. & Obama, T. The Oxidized Lipoproteins In Vivo: Its Diversity and Behavior in the Human Circulation. International Journal of Molecular Sciences vol. 24 Preprint at 10.3390/ijms24065747 (2023).

47. Tsujita, M., Melchior, J. T. & Yokoyama, S. Lipoprotein Particles in Cerebrospinal Fluid. *Arteriosclerosis*, Thrombosis, and Vascular Biology vol. 44 1042–1052 Preprint at 10.1161/ATVBAHA.123.318284 (2024).

48. Greenberg, M. E. et al. The lipid whisker model of the structure of oxidized cell membranes. Journal of Biological Chemistry 283, 2385–2396 (2008).

49. Spronck, E. A. et al. AAV5-miHTT Gene Therapy Demonstrates Sustained Huntingtin Lowering and Functional Improvement in Huntington Disease Mouse Models. Mol Ther Methods Clin Dev 13, 334–343 (2019).

50. Chen, H. J. & Mitchell, J. C. Mechanisms of tdp-43 proteinopathy onset and propagation. International Journal of Molecular Sciences vol. 22 Preprint at 10.3390/ijms22116004 (2021).

51. 51. Suk, T. R. & Rousseaux, M. W. C. The role of TDP-43 mislocalization in amyotrophic lateral sclerosis. Molecular Neurodegeneration vol. 15 Preprint at 10.1186/s13024-020-00397-1 (2020).

52. Healy, L. M., Stratton, J. A., Kuhlmann, T. & Antel, J. The role of glial cells in multiple sclerosis disease progression. Nature Reviews Neurology vol. 18 237–248 Preprint at 10.1038/s41582-022-00624-x (2022).

53. Wahrle, S. E. & Holtzman, D. M. Differential metabolism of ApoE isoforms in plasma and CSF. Exp Neurol 183, 4–6 (2003).

54. 54. Elliott, D. A., Weickert, C. S. & Garner, B. Apolipoproteins in the brain: Implications for neurological and psychiatric disorders. Clinical Lipidology vol. 5 555–573 Preprint at 10.2217/clp.10.37 (2010).

55. Bergmark, C. et al. A novel function of lipoprotein [a] as a preferential carrier of oxidized phospholipids in human plasma. J Lipid Res 49, 2230–2239 (2008).

56. Leibundgut, G. et al. Determinants of binding of oxidized phospholipids on apolipoprotein (a) and lipoprotein (a). J Lipid Res 54, 2815–2830 (2013).

57. Linton, M. F. et al. Phenotypes of Apolipoprotein B and Apolipoprotein E after Liver Transplantation. J. Clin. Invest. (1991).

58. Pitas, R. E., Boyles, J. K., Lee, S. H., Foss, D. & Mahley, R. W. Astrocytes synthesize apolipoprotein E and metabolize apolipoprotein E-containing lipoproteins. Biochimica et Biophysics Actcr 917, 148–161 (1987).

59. Zalocusky, K. A. et al. Neuronal ApoE upregulates MHC-I expression to drive selective neurodegeneration in Alzheimer’s disease. Nat Neurosci 24, 786–798 (2021).

60. Guo, J. L. et al. Decreased lipidated ApoE-receptor interactions confer protection against pathogenicity of ApoE and its lipid cargoes in lysosomes. Cell (2024) doi:10.1016/j.cell.2024.10.027.

61. Meneses, A. D. et al. APOE2 Exacerbates TDP-43 Related Toxicity in the Absence of Alzheimer Pathology. Ann Neurol 93, 830–843 (2023).

62. Bu, G. Apolipoprotein e and its receptors in Alzheimer’s disease: Pathways, pathogenesis and therapy. Nature Reviews Neuroscience vol. 10 333–344 Preprint at 10.1038/nrn2620 (2009).

63. Sjekloća, L. & Buratti, E. Conserved region of human TDP-43 is structurally similar to membrane binding protein FARP1 and protein chaperons BAG6 and CYP33. MicroPubl Biol (2024) doi:10.17912/micropub.biology.001388.

64. Barber, S. C. & Shaw, P. J. Oxidative stress in ALS: Key role in motor neuron injury and therapeutic target. Free Radic Biol Med 48, 629–641 (2010).

