## Supplementary figures and images for "AAV-Delivered Anti-PC-OxPL Antibody Fragments: A Novel Therapeutic Approach to Target ALS"

### Suppl Figures 1 - 3.pdf

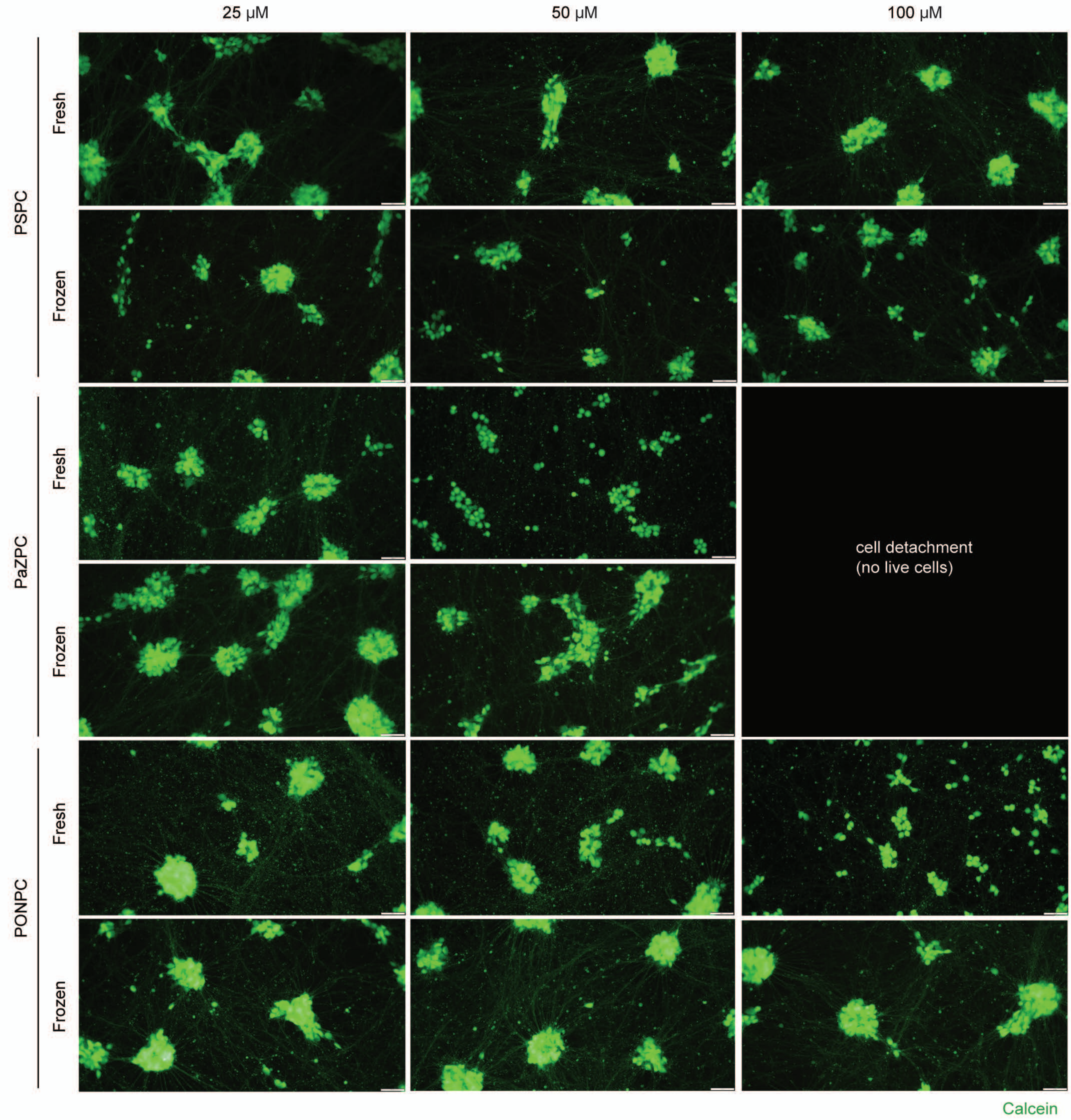

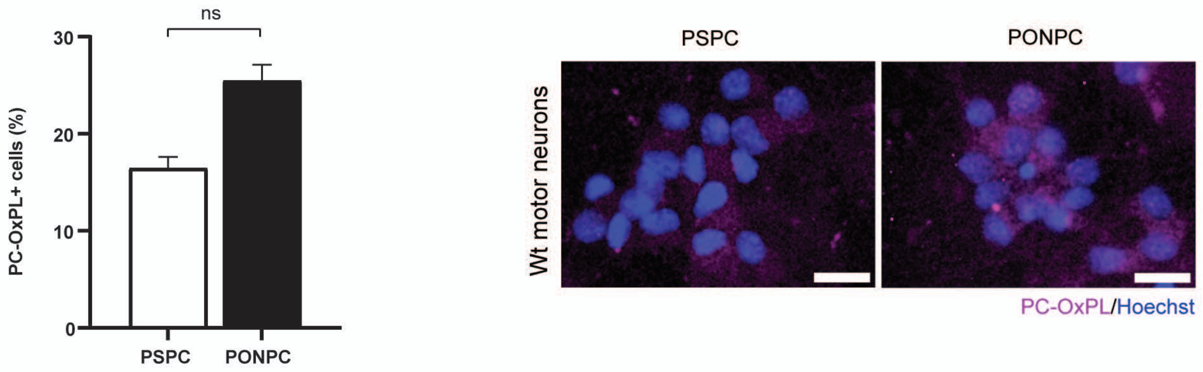

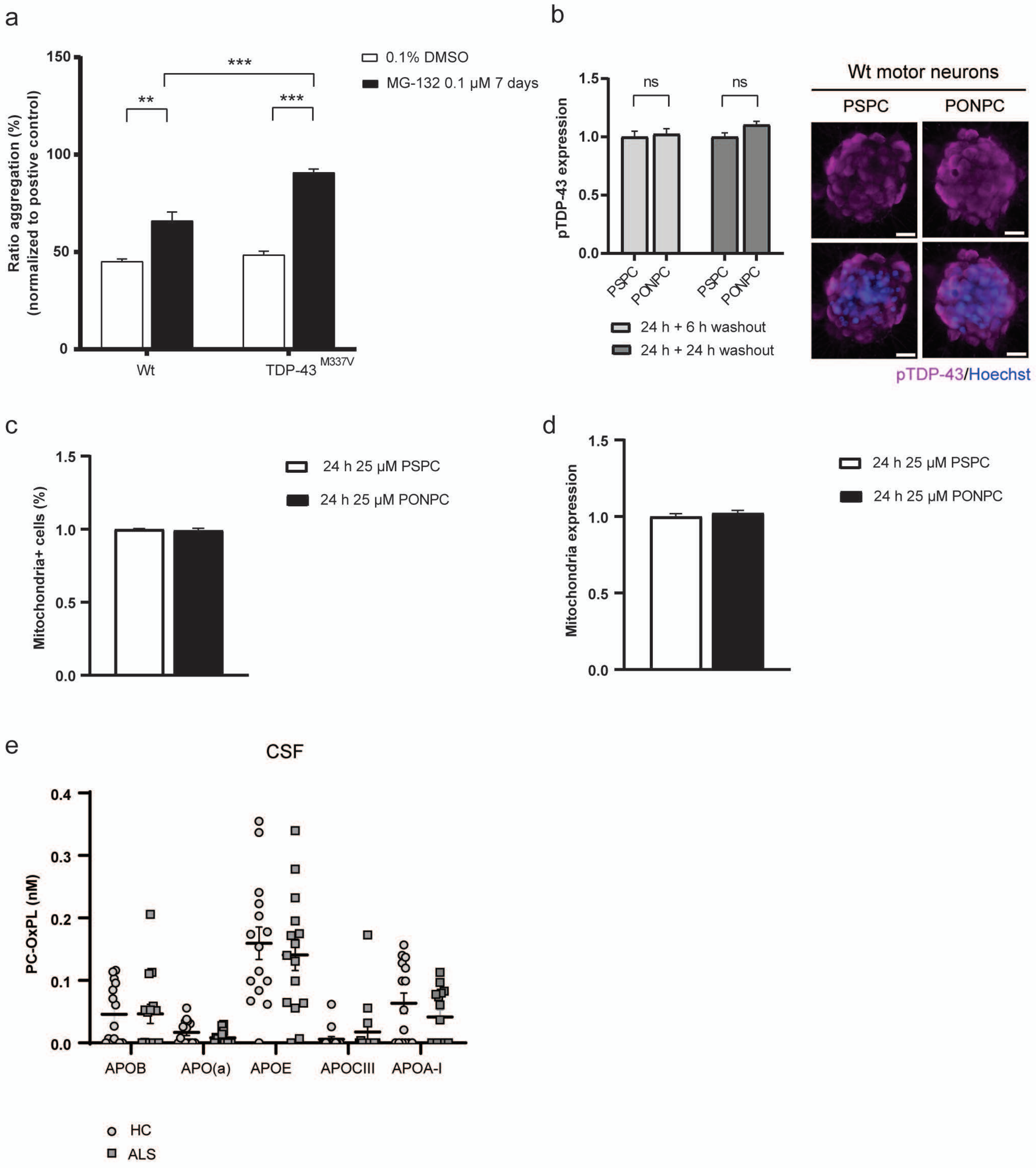

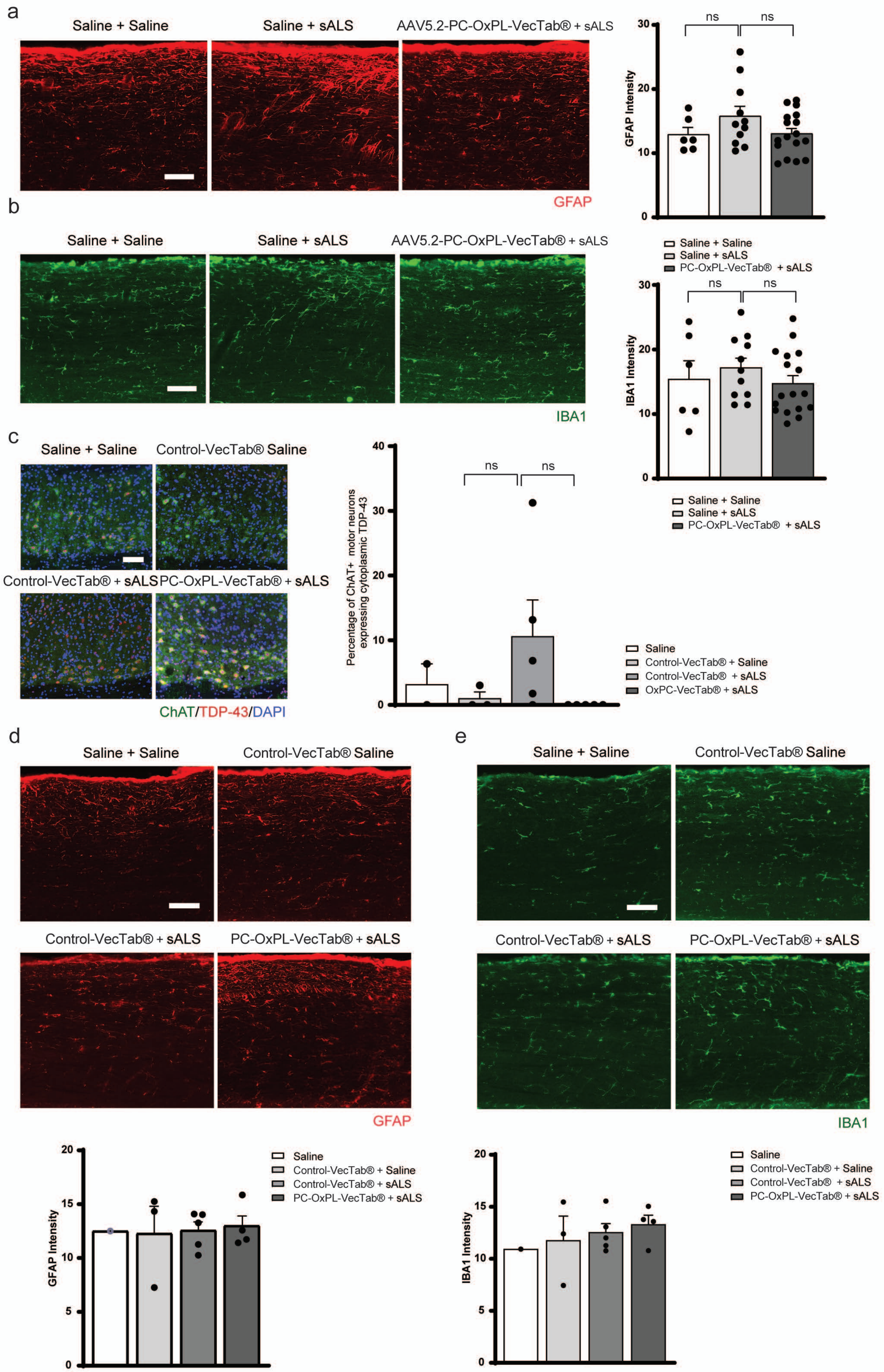
